# Projection-Specific Intersectional Optogenetics for Precise Excitation and Inhibition in the Marmoset Brain

**DOI:** 10.1101/2025.06.18.660378

**Authors:** Luke Shaw, Krishnan Padmanabhan, Amy Buckleaw, Jude F. Mitchell, Kuan Hong Wang

## Abstract

The primate cerebral cortex relies on long-range connections to integrate information across spatially distributed and functionally specialized areas, yet tools for selectively modulating these pathways remain limited. Here, we present an optimized intersectional viral and optogenetic strategy for precisely exciting and inhibiting projection-specific neurons in the common marmoset. Building on a mouse-to-marmoset pipeline, we first validated that optogenetic activation of inhibitory neurons (via AAV9-Dlx-ChR2) enables robust local cortical inhibition. We then combined retrograde delivery of Cre-recombinase (AAVretro-Cre) with locally injected Cre-dependent vectors encoding excitatory or inhibitory opsins (AAV8-FLEx-ChR2 or Jaws) to achieve directionally selective expression in callosal and frontoparietal pathways. Dual-opsin co-expression enabled precise stimulation or suppression of projection neurons in vivo with minimal off-target labeling. These results establish a scalable framework for projection-specific optogenetic interrogation of distributed circuits in primates, expanding the experimental toolkit for causal studies of higher-order brain function with enhanced anatomical precision and functional specificity.

**Graphical Abstract:** 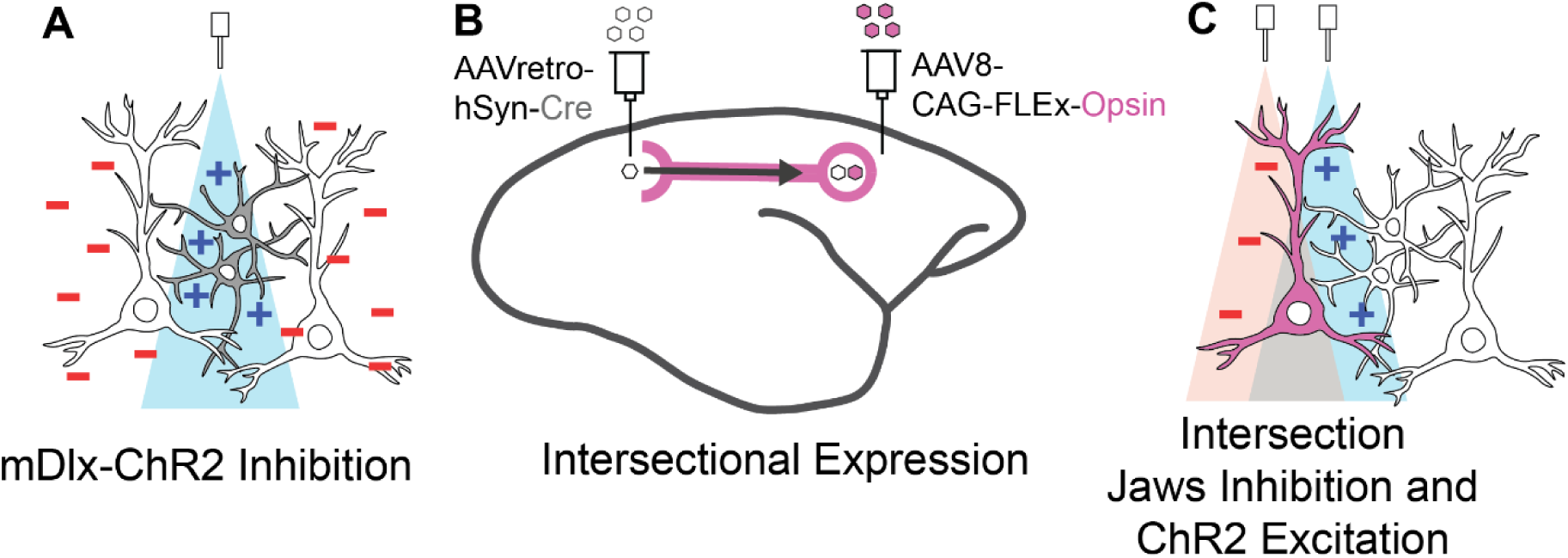

## INTRODUCTION

The coordinated activity of the cerebral cortex relies on long-range connections that integrate the neural activity of specialized processing regions^1,2^. A fundamental challenge of neuroscience is to decipher the architecture of these brain circuits and their functional impacts on neural activity and behavior. The advent of optogenetics, a technique that enables precise manipulation of neural firing through specific wavelengths of light, has significantly advanced our understanding of targeted circuits and neuronal populations^3^. However, the adaptation of this powerful toolkit for use in non-human primate (NHP) studies has progressed more slowly compared to its application in rodent models^4,5^. Given the close evolutionary proximity of NHPs to humans and their more complex cortical organization, further developing optogenetic tools in these species is essential for translational neuroscience.

Key features of NHP cortical architecture include larger brain sizes and extensive long-range neuronal projections that link functionally specialized cortical regions^2,6^. Compared to rodent neural architecture, NHP cortical areas are more functionally distinct and include elaborated intermediary processing regions, such as those in the frontal and parietal association cortices^7–9^. The common marmoset (*Callithrix jacchus*), a small, lissencephalic New World primate, is particularly valuable for translational neuroscience due to its cortical architecture, which closely resembles that of Old-World monkeys and apes while offering greater experimental accessibility and animal availability^10,11^. These advantages make the marmoset an ideal platform for adapting and scaling up projection-specific optogenetic strategies initially developed in rodents.

Selective targeting of neural circuit components has been widely enabled in rodent models through the development of transgenic Cre driver lines^12,13^. In contrast, comparable strategies in NHPs remain limited due to the technical and logistical challenges of establishing stable transgenic Cre lines^4,14,15^. While adeno-associated virus (AAV)-based delivery of opsins reliably drives expression in NHPs^16–18^, targeting specific cell classes remains difficult. Most studies rely on pan-neuronal promoters to ensure high expression levels, but this broad targeting simultaneously activates both excitatory and inhibitory neurons, often producing ambiguous or weak net effects on local circuit activity ^4,5,19^. To limit the scope of expression, an inhibitory neuron-specific Dlx promoter has been used to drive ChR2 expression via AAV delivery, producing reliable local cortical inactivation in both rodents and macaque monkeys^20–24^. Nevertheless, because this strategy inhibits many local neurons regardless of projection identity, it does not permit pathway-specific circuit dissection. Achieving this level of specificity requires methods that restrict expression to projection-defined neuronal populations.

Another method that has shown promise in NHPs is targeting projection pathways. For example, local optical stimulation of ChR2-labeled axons from frontal cortex to superior colliculus reliably evoked eye movements^25^. However, effective applications of this approach are largely limited to pathways without reciprocal connections, as many AAV serotypes undergo both anterograde and retrograde transports, albeit with varying efficiencies ^18,26–29^. This raises the risk of unintended expression in neurons projecting back to the injection site, which can confound circuit-specific manipulations. In addition, although axon terminal inhibition via optogenetics is feasible, its efficacy is constrained by the biophysical properties of axons, their restricted optical accessibility, and the need for anatomically precise targeting to ensure pathway specificity^30–33^. Together, these limitations highlight the need for strategies that enable directionally restricted expression in anatomically defined projection pathways, while also supporting bidirectional modulation, either excitation or inhibition, of the targeted neuronal population.

To overcome the limitations in current NHP optogenetic approaches, we developed an intersectional strategy that leverages viral transport across long-range axonal projections. This approach, widely established in rodent optogenetic studies^27,34–36^ and piloted in a few macaque anatomical and behavioral studies^26,37^, involves injecting a Cre-encoding virus engineered for retrograde transport at the projection target while simultaneously injecting a Cre-dependent opsin construct at the projection origin. A key challenge in this intersectional method is identifying viral vector pairs that work synergistically to achieve high labeling specificity. Different AAV serotypes have distinct cell tropisms and transport directions (retrograde or anterograde), which must be carefully matched^18,29,38^. While AAVretro is effective for retrograde transport at projection terminals^27^, the complementary virus at the soma site must exhibit minimal retrograde affinity to reduce off-target or “leaky” expression in reciprocal pathways. This is a particular concern in primate cortex, where bi-directional connectivity is common^6,18,39^. Furthermore, maintaining healthy and stable expression requires careful optimization of viral serotypes and particle concentrations^40,41^, a process complicated by the limited availability of NHP subjects. These constraints underscore the importance of an optimized, intersectional viral toolkit tailored for primate use.

In this study, we first validated local optogenetic inhibition in marmoset cortex using AAV9-mDlx-ChR2 and surface light delivery, confirming robust suppression of cortical activity across cortical depth using laminar array electrophysiological recordings. Building on this foundation, we next developed and optimized a projection-specific, dual-opsin intersectional strategy to enable selective excitation and inhibition of anatomically defined cortico-cortical pathways. Following successful testing and titration in mouse models, we translated this strategy to the marmoset, using an AAVretro/AAV8 combination that maximized projection specificity while minimizing leaky expression and tissue over-transduction. We further refined this approach to achieve co-expression of the excitatory opsin ChR2 and the inhibitory opsin Jaws in the same projection-defined population within both callosal and frontoparietal pathways of the marmoset cortex. This dual-opsin configuration enabled bidirectional control of neural activity in projection-specific populations, validated by in vivo electrophysiology, and establishes a scalable and effective platform for projection-specific, bidirectional optogenetic interrogation of distributed cortical circuits in NHPs.

## RESULTS

### mDLX-ChR2-mediated Local Inhibition in the Marmoset Brain

To first validate our optogenetic preparation in marmoset cortex, we tested the use of mDlx-ChR2 for suppressing local cortical populations, a method that has proven robust in other NHP preparations over recent years^20,21^. The use of mDlx-ChR2 offers area-specific control of cortical inhibition with millisecond response timescales and minimal invasiveness. To examine the labelling patterns and optogenetic effects of virally delivered mDlx-ChR2 in the marmoset, we injected AAV9-mDlx-ChR2 into the premotor cortex of a marmoset. After an expression period, we performed electrophysiological recordings while the animal was head-fixed, alert, and engaged in viewing natural images (**Figure 1A**). Viral micro-injections here and in the following tests were delivered at 2-3 sites in a 3 mm x 3 mm craniotomy to maximize expression coverage over an area of cortex. Histological analysis confirmed robust labeling of mDlx-ChR2+ cells with inhibitory interneuron morphology (**Figure 1 B,C**). In the premotor cortex section with the highest amount of labeling, a total of 288 ChR2+ cells were identified. In this coronal section, labeled somas and dendrites extended over an area of 3.67 mm^2^, with a soma density of 78.47 cells/mm^2^ at the center of expression. Additional coronal sections located 0.5 mm anterior and 0.5mm posterior to this section were also quantified. These sections showed labeled cortical areas of 3.43 mm^2^ and 3.52 mm^2^, with corresponding soma densities of 74.34 cells/mm^2^ and 73.58 cells/mm^2^, respectively. The area of expression extended approximately 1.6 mm anterior and posterior to the central section before tapering off. This extent of labeling likely reflects the spatial separation between injection sites within the craniotomy.

**Figure 1.**
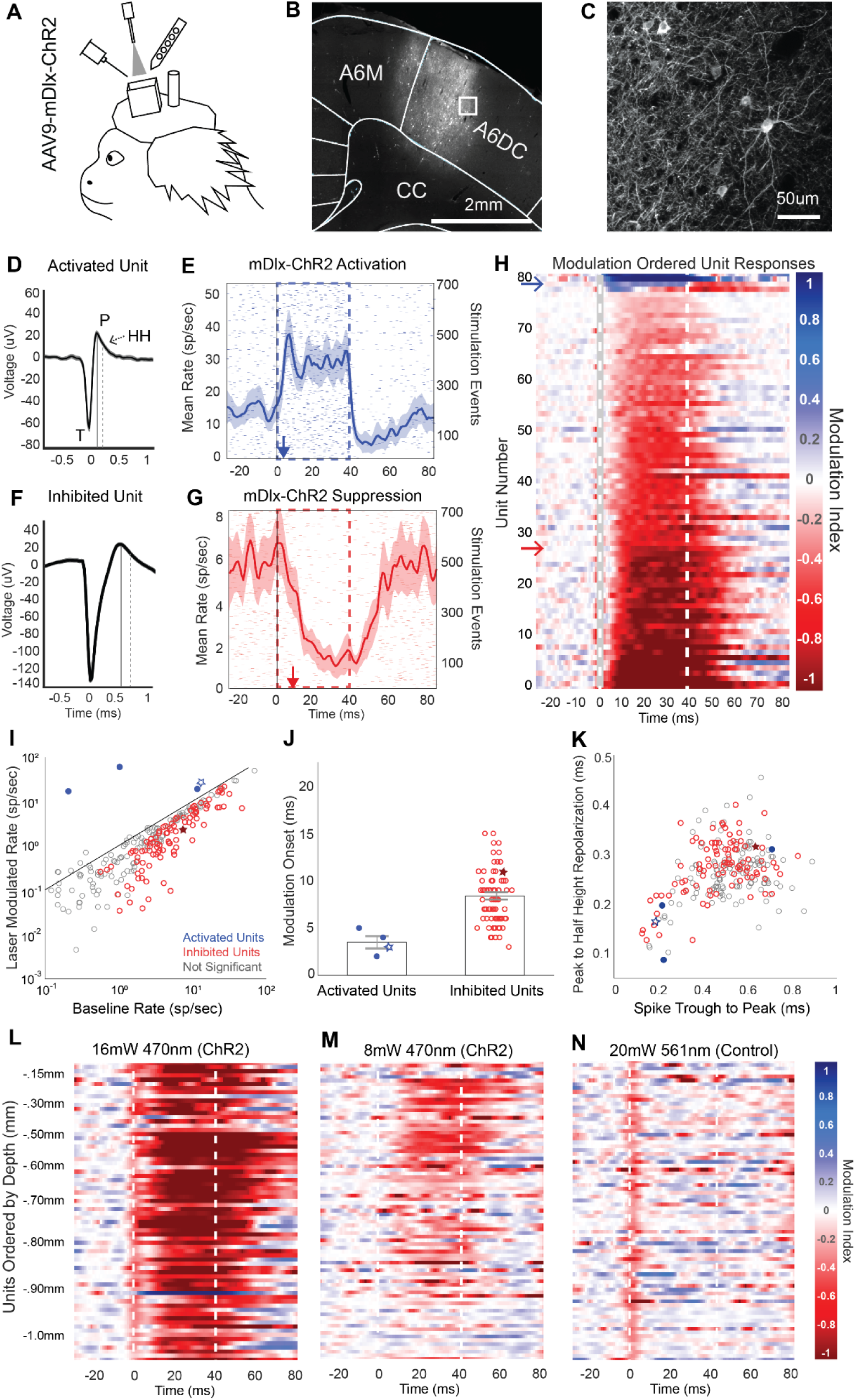
AAV9-mDlx-ChR2-mediated Local Cortical Inhibition in the Marmoset. **A** Electrophysiology during optical stimulation was conducted in an awake, head fixed marmoset freely viewing natural images. AAV9-mDlx-ChR2 was injected within a craniotomy over premotor area A6DC. **B-C** Native fluorescence of mDlx-ChR2+ inhibitory interneurons. **D** Average spike waveform of an excited unit. T indicates the trough of the spike. P indicates the post-depolarization voltage peak, marked in time by the solid line. HH indicates the half height of the repolarization voltage curve, which is marked in time by the dashed line. This unit is displayed as a red star in panels I-J. **E** Laser triggered PSTH of the unit in D. The red arrow indicates the modulation onset time. Shaded regions indicate ±2 standard errors of the mean (SEM). **F** Average spike waveform of an example suppressed unit. The solid and dashed lines indicate the peak and repolarization half height, as in D. This unit is displayed as a blue star in panels I-J. **G** Light triggered PSTH of the unit in F. The blue arrow indicates the modulation onset time. **H** Spike-sorted units ordered by degree of laser-induced firing modulation. Red indicates excitation; blue indicates suppression. Dotted lines indicate the laser stimulation period. Modulation index calculation is detailed in Methods. **I** Laser-triggered firing rate of units plotted against their baseline firing rate. Red points are significantly activated relative to baseline; blue points are suppressed. **J** Excited and inhibited units plotted by the onset of laser-triggered modulation. Onset was estimated by the first time point at which the laser-modulated firing rate rose above or fell below baseline ±2 SEM. **K** Spike waveforms characterized by trough (T) to peak (P) duration and peak (P) to repolarization half-height (HH) duration. **L-N** Blue laser (470 nm) power was reduced to test the modulation of laser-triggered optogenetic effect in a continuous recording. High-power green light (561 nm) was used as a control to confirm no neural suppression from laser-induced heating. Units are ordered by cortical depth. Apparent suppression at time 0 ms reflects software-based removal of spikes coinciding with the optical-electric artifact triggered by laser onset.

ChR2 delivery using the mDlx promoter has been shown to specifically target inhibitory interneurons^21^. Optogenetic stimulation using this construct stimulates inhibitory cells, which act to suppress local network activity due to their local inhibitory projections. Electrophysiological signatures of this stimulation should thus show two general trends: a small population of activated units, putative inhibitory mDlx+ labeled neurons that increase their firing rate under ChR2 stimulation, and a more sizeable population of inhibited units whose firing rate drops off due to the activity of the excited inhibitory interneurons.

We tested if our preparation for recording and optogenetic stimulation would be sufficient to produce robust suppressive effects. We delivered light at the cortical surface through a thin layer of silastic protecting the underlying brain tissue. Spike sorted data from five recording experiments at different sites within the craniotomy were combined and analyzed as a function of light-triggered spike rate modulation and cortical depth. These experiments delivered 40 ms pulses of ChR2 light stimulation at a rate of 5 Hz in 30 second runs (16 mW, 470 nm: 127.32 mW/mm^2^ at fiberoptic tip, estimated 38.2 mW/mm^2^ at cortical surface). Over five recordings, a total of 279 unique units were identified with well isolated spike waveforms (see Methods). Eighty-one of these units showed modulation during the light activation period, with firing rates deviating above or below baseline by more than two times the standard error of the mean (SEM).

A subset of neurons showed excitation on ChR2 stimulation, consistent with characteristics of inhibitory interneurons, while the majority of neurons were suppressed. We examined the latency and direction of light triggered modulation timing as well as spike waveform characteristics of modulated units to determine whether they match the expected features of directly activated inhibitory cells or inhibited downstream cells. Due to the small size and short refractory period of typical inhibitory interneurons, we expect that a directly activated inhibitory cell should have spike waveform with short trough (T) to peak (P) duration as well as short peak to refractory half height (HH) duration^42,43^. A unit with such characteristics (**Figure 1D**) was identified, which also shows short latency activation within the laser-on period (**Figure 1E**). This unit’s spiking activity first deviated from baseline 3 ms post-laser onset, which is typical of direct ChR2-mediated activation^3^. Further exploration of the population also showed units with spike waveforms of longer duration and longer half height refractory periods, typical of pyramidal cells (example, **Figure 1F**)^42,43^. One such broad spiking neuron was inhibited by light stimulation and had a slower latency in its suppression (**Figure 1G**). The onset of this effect was 11 ms, which follows from it being an indirect effect mediated by direct activation of an inhibitory neuron.

We found that the majority of cells showed a general trend of activity suppression in response to mDLX-ChR2 stimulation, while only a small subset were excited (**Figure 1H**). To illustrate this activity modulation on a population scale, identified units were grouped as excited or inhibited using a modulation index. This index was calculated by subtracting the baseline firing rate from the instantaneous firing rate (binned at one-millisecond resolution) and dividing the result by the sum of the instantaneous and baseline rates. Of the 279 units, 77 showed significant negative modulation and 4 showed significant positive modulation (**Figure 1I**). For this classification, we looked specifically at an early window of 10 ms to 25 ms to disregard slower network effects. As expected, when we separated the population of activated cells from the population of inhibited cells and examined the modulation onset times for these groups, the small subset of activated cells were significantly modulated within 5 ms (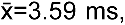 σ=.42), while inhibited cells were significantly modulated at slower latencies in a range of 3 to 15 ms (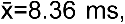 σ=.29) (**Figure 1J**). Analyses of spike waveforms demonstrated that excited units generally demonstrated characteristics of inhibitory interneurons, including narrow spike widths and short repolarization durations. When spiking waveform trough to peak durations were plotted against peak to half height repolarization durations, three of the four activated units appeared within a cluster of short duration waveform features whereas the suppressed units appeared within a larger cluster with longer duration spiking features (**Figure 1K**).

In addition, we found that light modulation impacted neurons across cortical depths even though the optical fiber was placed above the silastic protecting the cortical surface. Of the activated units, the anatomically deepest unit was found at 1.45 mm relative to the cortical surface. Of the inhibited units, the anatomically deepest unit was found at 1.98 mm relative to the cortical surface, which is approximately the bottom extent of A6DC cortex.

In a separate control recording, we performed tests of laser power and wavelength to validate that suppression effects were not due to tissue heating and to further confirm that deep cortical layers could be modulated by light at the surface. First, a laser power reduction test confirmed that optogenetic modulation is concomitantly reduced. A total of 71 units were found in this series of recordings and are shown as heatmaps arranged by their depth relative to the cortical surface (**Figure 1L-M**). Using 16 mW of blue light (470 nm, 127.32 mW/mm^2^ at the tip), we observed consistent suppression of neuronal responses at depths up to 1.0 mm. In contrast, a lower power of 8 mW (470 nm, 63.66 mW/mm^2^ at the tip) produced more restricted effects, limited to a depth range of approximately 0.30 to 0.80 mm. As a wavelength control, we applied a 20 mw green laser (561 nm, 159.15 mW/mm^2^), which should minimally activate ChR2 while inducing comparable tissue heating, and found no neural effects at any depth (**Figure 1N**). Apparent suppression at time 0 ms reflects software-based removal of spikes coinciding with the optical-electric artifact triggered by laser onset. Additional tests were performed using 4 mW blue light stimulation (470 nm, 31.83 mW/mm^2^ at tip), which showed almost no suppression (**Supplemental Figure 1**). For all conditions, a mean modulation score across units in the 10 ms to 40 ms post laser onset window was significant at 16, 8 and 4 mW blue light (two-tailed student’s t-tests, 16 mW: µ =-.73, p=8.3E-30; 8 mW: µ =-.23, p=8.4E-16; 4 mW: µ=-.10,p=6.5E-8), while the green light control and shuffled null control were non-significant as expected (Green: µ =.03, p=.086; Null: µ = -.02, p=.23). Finally, this experiment confirmed that in the higher power condition (16 mW), stimulation from the cortical surface was sufficient to suppress activity even at the deepest sites at slight over 1 mm of depth (**Figure 1L**).

### AAV9-based Intersectional ChR2 Expression Lacks Specificity

To first test the efficacy of projection-direction-specific intersectional labelling in the marmoset, we utilized an AAV9 serotype to deliver CAG-FLEx-ChR2 to the cell bodies of cross-callosal projection cells in the right hemisphere of the premotor cortex, combined with AAVretro-hSyn-Cre injection in the left hemisphere (**Figure 2A**). AAV9 was our first choice due to the high efficiency of this serotype, which has been documented in marmoset cortex^7,18,30^. Left and right premotor cortex are connected by dense, bi-directional callosal projections ^6,44^ and thus this circuit served as an ideal ground for tests of a robust and relatively short-range projection.

**Figure 2.**
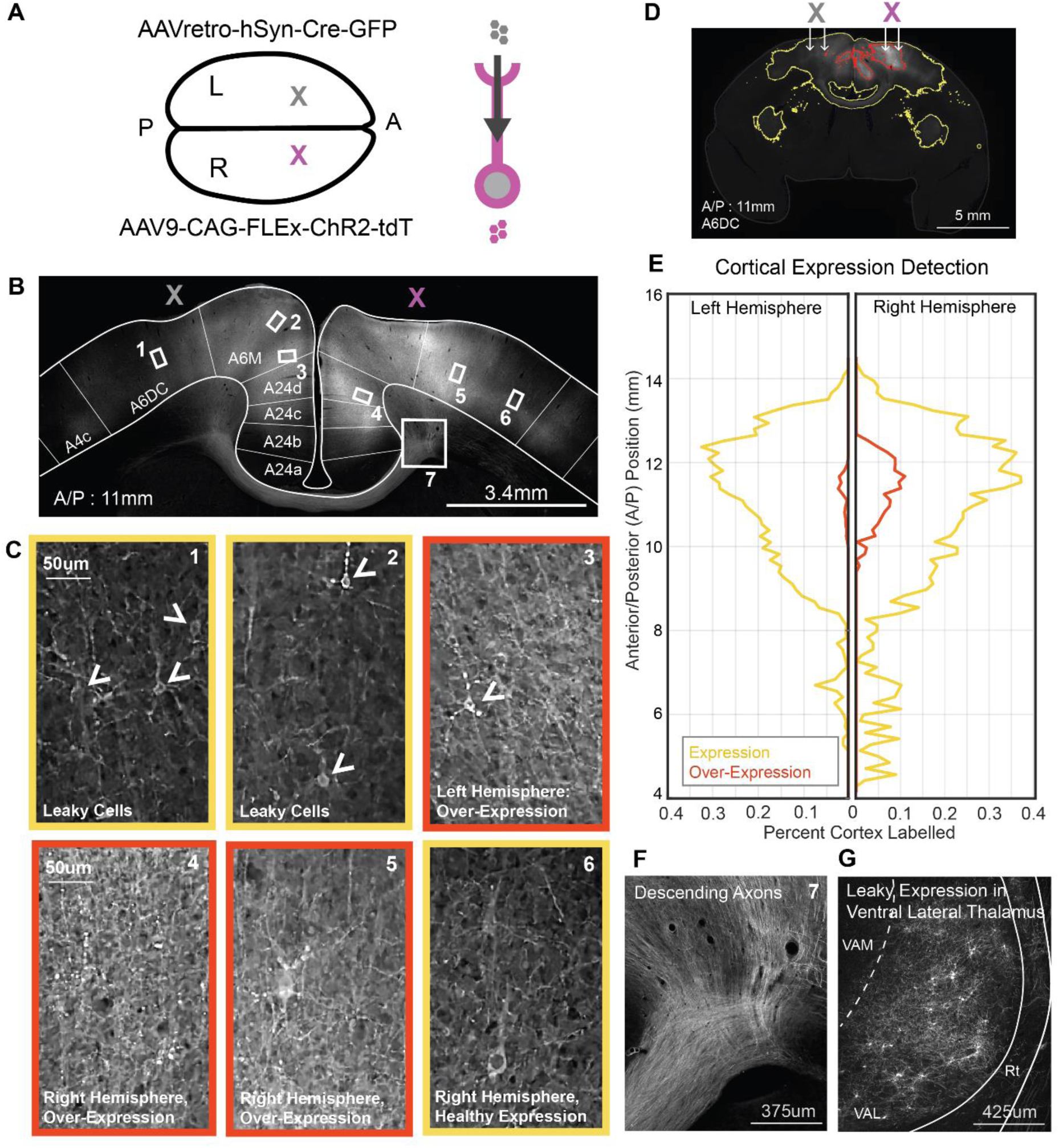
AAV9-based Intersectional Strategy Yields Efficient but Leaky Cross-Callosal ChR2 Expression. **A** Intersectional viruses were delivered to left (Cre-gray) and right (ChR2-magenta) A6DC in marmoset premotor cortex. **B** Example native fluorescence of tdTomato in premotor cortex at the section showing peak expression, with numbered boxes for subsequent zoom-ins. **C** Example cortical ROIs at the Cre injection site and ChR2 injection site showing native tdTomato fluorescence. Leaky expression is visible at the Cre site (arrowheads), with variable expression levels across cases. Panels with red borders highlight examples of over-expression, characterized by blebbing puncta and elevated parenchymal fluorescence. Panels outlined in yellow indicate typical expression patterns. **D** An example coronal section shows results of automated tissue classification based on native tdTomato fluorescence. Despite visible subcortical axonal labeling in the basal ganglia and claustrum, only cortical regions were included in quantification. **E** Both hemispheres showed widespread cortical labeling, quantified as the percent of cortical area exhibiting either healthy expression or over-expression relative to total cortical area in each coronal section. **F** Labeled decussating axons highlight the robustness of cross-callosal tdTomato expression. **G** “Leaky” somatic labeling by tdTomato was detected in motor-related thalamic regions, including Ventral Anterior Lateral Thalamus (VAL). VAM: Ventral Anterior Medial Thalamus; Rt: Reticular nucleus.

Initial findings suggest that intersectional labeling with AAV9 yields highly efficient expression. This approach produced a generally symmetric labeling pattern across the cerebral hemispheres, with cross-callosal projections terminating in a columnar, patchy manner within bilaterally corresponding regions (**Figure 2B; Supplemental Figure 2**). However, dense parenchymal labeling made cell quantification challenging. Although labeling in the left hemisphere would be expected if limited to axonal terminals, the presence of numerous labeled cell bodies (**Figure 2C1–2**) suggests reduced specificity and possible leakiness into the reciprocal projection pathway. Furthermore, some cortical areas showed signs of overexpression, evidenced by the accumulation of labeled puncta in dendritic arbors and somas (**Figure 2C3–5**), whereas other regions displayed healthy expression characterized by uniform membrane labeling (**Figure 2C1, 2, 6**).

To quantify the extent of expression in each hemisphere, we used an automated tissue classification approach based on the tunable artificial neural network (ANN) supplied in QuPath software. This approach enabled classification of epifluorescence images across the entire sectioned brain, with sampling at one section every 300 um. The ANN was trained to discriminate healthy expression (yellow), over-expression expression (red), and un-labelled tissue (**Figure 2D**). Training sets included examples of labelled axon tracks, terminal fields, and cell bodies. Notably, the extent of labelling, shown as the percentage of tissue area covered by labels, was widespread in both hemispheres with higher incidences of overexpression in the right hemisphere (**Figure 2E**). Regions of overexpression in the right hemisphere spread 4 millimeters over the anterior/posterior coordinate frame and are also detected in the left hemisphere close to the injection site.

AAV9’s high transduction efficiency and endogenous retrograde transport properties^18,38,45^ contributed to off-target labelling not only at the Cre injection site in the left cortex, but also in the thalamus. While labeling of descending cortical projections (**Figure 2F**) and terminal fields in subcortical structures (**Figure 2D**) was expected, we also observed extensive cell body labelling in the right Ventral Anterior Lateral (VAL) thalamus (**Figure 2G**), a motor-related region known to project to the premotor cortex. This unexpected expression pattern may result from retrograde transport of both AAVretro and AAV9 in VAL-originating projections that ascend to the right premotor cortex and decussate to the left premotor cortex via the corpus callosum. Similar decussating thalamo-cortical labelling patterns have been reported in frontal cortical areas of macaques^46^, as well as in mice^47^ and rats^48^. While informative, such off-target labeling compromises projection specificity. To address this, we tested alternate AAV serotypes in the mouse to further improve labeling precision.

### Testing Intersectional ChR2 and Jaws Co-Expression using AAV8 in the Mouse

A wide range of AAV serotypes, promoters, and viral titers can be used in combination with AAVretro to achieve intersectional labeling of a single projection direction, though their efficacy and specificity can vary considerably. Most studies using these tools in mice^18,49–51^ employ significant numbers of test animals to pilot different serotype, promoter, and titer combinations to optimize the desired expression properties. However, in non-human primates the numbers of animals for use can often be limited. This challenge for viral optimization became apparent in our first test in marmoset cortex, detailed above, where we found AAV9 led to overexpression and leaky expression in the reciprocal projection pathway. Our findings highlight the need to optimize viral parameters using multiple test animals, ultimately including mice as well as marmosets.

To achieve more specific labelling of targeted projections, we tested in the mouse an AAV8 serotype with less retrograde affinity in an effort to eliminate leaky expression in the reciprocal pathway to obtain better directional specificity. In addition, we tested the feasibility of including not only the excitatory opsin ChR2 but also of the inhibitory optogenetic chloride pump Jaws to suppress the projection pathway^52^. AAV8 was selected because comparative analyses of AAV serotype expression profiles indicated that AAV8 demonstrates a more balanced cortical expression profile, providing efficient local transduction while avoiding excessive retrograde transport^27,38^. We again used a cross-callosal injection strategy, delivering AAVretro-hSyn-Cre-BFP into the left M1/M2 region and a 1:1 mixture of AAV8-CAG-FLEx-ChR2-tdTomato and AAV8-CAG-FLEx-Jaws-GFP into the right M1/M2 (**Figure 3A**).

**Figure 3.**
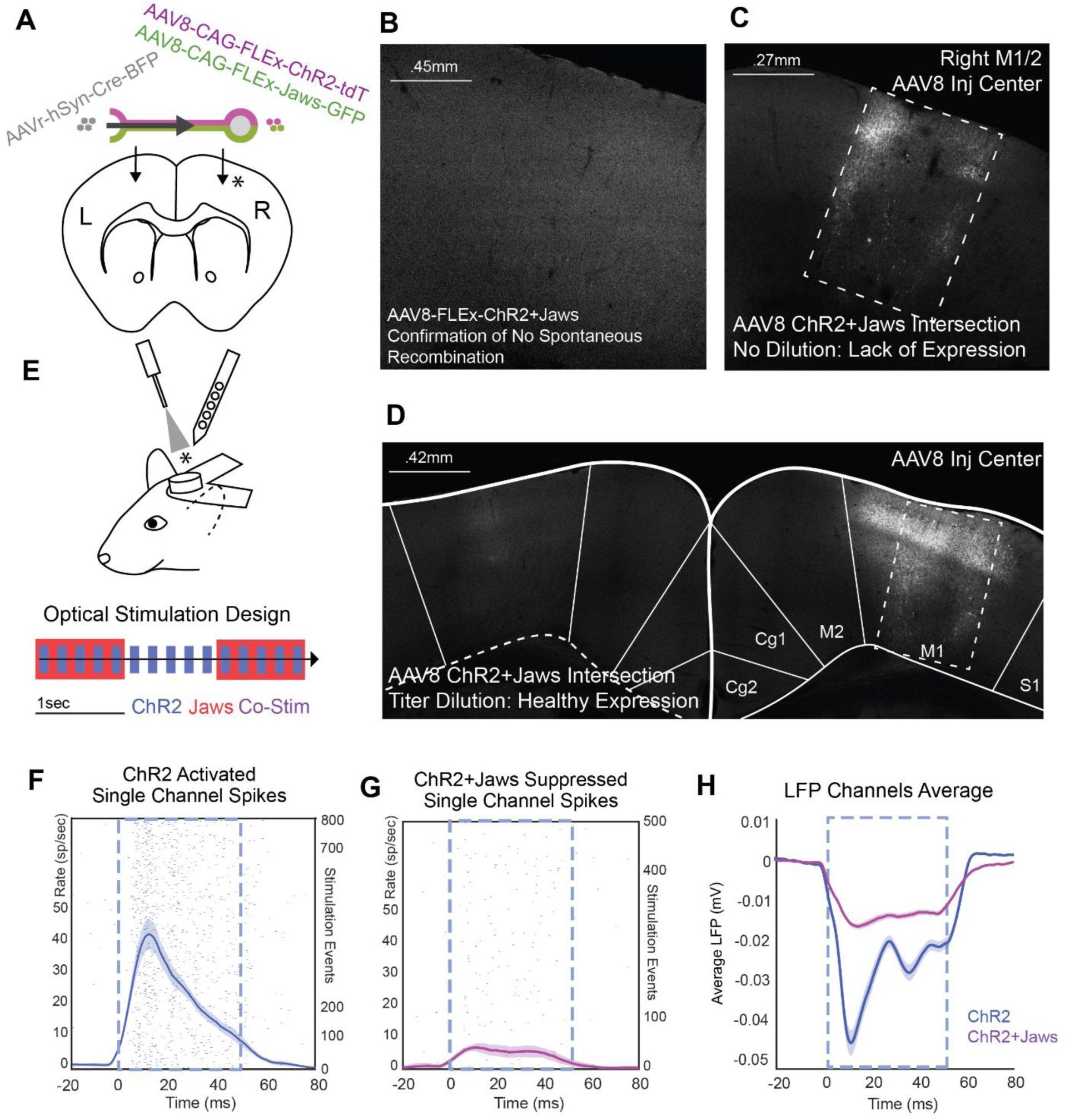
Evaluation and Titration of AAV8-Based Intersectional Jaws and ChR2 Expression in the Mouse Brain. **A** Design of a cross-callosal intersection strategy in the mouse using a 1:1 mixture of AAV8-FLEx-ChR2-tdTomato and AAV8-FLEx-Jaws-GFP injected into one hemisphere, and AAVretro-hSyn-Cre injected into the contralateral hemisphere. **B** Injection of AAV8-flex-ChR2+Jaws in the absence of Cre virus confirmed no unexpected recombination-related expression. This and subsequent panels present a grayscale composite of native tdTomato (ChR2) and GFP (Jaws) fluorescence signals. **C** Undiluted viral solutions produced poor expression around the injection site (dashed box), which could not be attributed to mechanical damage from the injection. **D** A 1:2 dilution of the AAV8-ChR2+Jaws mixture yielded healthy expression near the injection site (dashed box). Axon terminals were evident in the contralateral hemisphere at the AAVretro-Cre injection site. **E** Optical stimulation and electrophysiological recording design in head-fixed, awake mice. ChR2 stimulation pulses (blue) were delivered with or without concurrent long-duration Jaws pulses (red). Co-stimulation events are shown in purple. Recordings were obtained at the site of ChR2+Jaws injection. **F** ChR2 laser stimulation induced an increase in firing rate, as shown in the PSTH of multi-unit activity from a single recording channel (± 2 SEM). **G** During co-stimulation, the ChR2-driven excitation was effectively suppressed by Jaws stimulation. **H** This effect was evident in the grand average of local field potentials across multiple recording channels.

One advantage in using mice for these tests is that it is possible to perform a larger number of controls with test animals. In total we used 26 mice to optimize this strategy with AAV8, to test alternate serotypes, and to validate optogenetic stimulation in electrophysiology. We first performed a test using the AAV8 vectors to confirm there was no expression of ChR2 in the absence of Cre, which might occur due to spontaneous recombination reverting FLEx-ChR2. This was performed using a 1:1 mixture of the AAV8-FLEx-ChR2/Jaws vectors injected into the M1 cortex. Our tests indeed showed no identified Cre-independent expression from the AAV8 vectors (**Figure 3B**).

We also examined if poor tissue health and over-expression could be mitigated by adjusting the dilution of the AAV solution. AAV injection-related cytotoxicity can result from excessively high viral titers or residual cytotoxins from the production process. Diluting the viral solution effectively reduces the risk associated with both factors. We first tested a fresh batch of virus without dilution. A total of 400 nL of a 1:1 mixture of AAV8-CAG-FLEx-ChR2 (3E13 gc/mL) and AAV8-CAG-FLEx-Jaws (5E13 gc/mL) was injected into the right M1 cortex, while AAVretro-hSyn-Cre (1E14 gc/mL) was injected into the contralateral M1 cortex. The virus injection site showed poor tissue health and a lack of expression at the center (**Figure 3C**, center of dashed rectangle), with stronger expression observed in the surrounding tissue. This pattern prompted us to perform dilution experiments. We found that a 1:1 mixture of AAV8-CAG-FLEx-ChR2 diluted 1:2 (final titer: 1E13 gc/mL) and AAV8-CAG-FLEx-Jaws diluted 1:5 (final titer: 1E13 gc/mL) was sufficient to prevent cytotoxicity while maintaining adequate expression levels (**Figure 3D, right hemisphere)**.

Having optimized the viral dilution, we performed head-fixed, awake electrophysiological recordings to test if co-expression of Jaws and ChR2 enabled robust suppression and excitation of the callosal projection. To test the effects of ChR2 stimulation and combined ChR2 and Jaws stimulation, we delivered trains of 5Hz, 50 ms pulses using a 470 nm laser (159.15 mW/mm^2^ at the tip) for ChR2 activation, concurrently with overlapping 0.5 Hz, 1-second pulses of 561 nm light (159.15 mW/mm^2^ at the tip) for Jaws activation (**Figure 3E**). Optical stimulation was delivered by a fiber-optic cannula placed directly above the brain surface and recordings were performed with a 64-channel multielectrode array at the AAV8 injection site where the cell bodies of projection neurons were labeled. Neural spiking activity was identified on each recording channel and peri-stimulus time histograms were generated for laser pulse events. A clear time-locked increase in firing rate was observed in response to ChR2 stimulation events in an example unit (**Figure 3F**). However, in the same unit, this response was greatly reduced when ChR2 stimulation events occurred during Jaws stimulation events (**Figure 3G**). These effects were also evident in local field potential (LFP) recordings across the population of channels, reflected by a reduction in the mean negative deflection (**Figure 3H**, shown with 1x SEM error fields). The peak voltage deflection induced by ChR2 stimulation was reduced by 72.1% during concurrent Jaws stimulation. These findings indicate both the co-expression of Jaws and ChR2 in recorded units and the ability of Jaws stimulation to suppress ChR2 driven neural spiking activity. Having demonstrated efficient and specific intersectional co-expression of ChR2 and Jaws using AAV8 in the mouse, we next translated this approach to the marmoset.

### Intersectional AAV8-Mediated ChR2 and Jaws Co-Expression in the Marmoset Premotor Callosal Projections

Applying AAV8 for intersectional co-expression of ChR2 and Jaws in marmoset cortex resulted in greater specificity than AAV9 for targeting the cross-callosal projection. Using dilutions established from mouse testing, we injected AAVretro-hSyn-Cre-BFP into the left premotor/motor cortex (A6DC/A4) and a 1:1 mixture of AAV8-CAG-FLEx-ChR2-tdTomato and AAV8-CAG-FLEx-Jaws-GFP into the right premotor/motor cortex (**Figure 4A**). This approach produced robust and widespread expression at the AAV8 injection site, spanning approximately 2.5 mm mediolaterally at the peak (**Figure 4B,C**), with labeling in both superficial and deep layers consistent with the distribution of callosal projection neurons. The observed patchy expression pattern may reflect a combination of the injection strategy, which involved two separate 1.5 μL injections per hemisphere at two depths, and the specific connectivity of the intersected projection. Tissue disruption from subsequent electrophysiological recordings may also contribute to this pattern (**Supplemental Figure 3**). Importantly, unlike prior high-titer AAV8 injections (**Figure 3C**), this expression was not associated with a loss of healthy neurons within the injection site. Morphological analysis confirmed healthy cell bodies and dendritic arbors with clear co-expression of ChR2 and Jaws (**Figure 4D**, circles).

**Figure 4.**
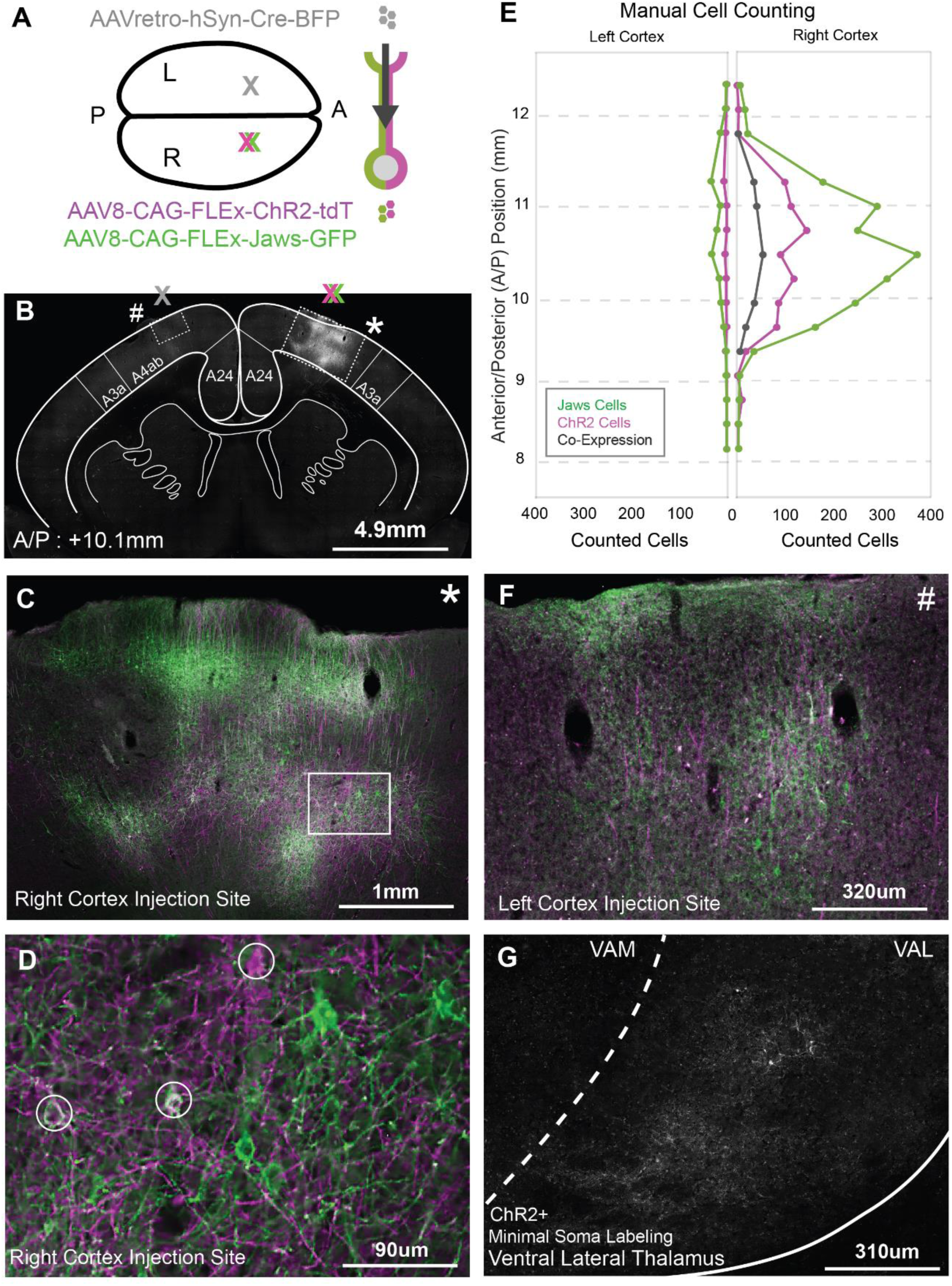
Efficient and Specific Cross-Callosal Intersectional Expression of ChR2 and Jaws using AAV8. **A** Intersection viruses were tested in the marmoset targeting the callosal projection from right A6DC/A4ab to left A6DC/A4ab. **B** Overview of the injection site in a coronal section, displaying a grayscale composite of native fluorescence of ChR2-tdTomato and Jaws-GFP. Squares indicate zoom-ins for panels C and F. **C** Zoom-in of the AAV8-ChR2 (magenta) + Jaws (green) injection site shows superficial and deep cortical layer expression. **D** Higher magnification from Panel C shows healthy soma and dendritic labelling. Circles mark neurons co-expression Jaws (green) and ChR2 (magenta). **E** Manual counts of labeled somata in right and left cortex show greater expression in the right hemisphere, indicating low leakiness. Each dot represents the number of labeled somata in a single coronal section. **F** Zoom-in of the Cre injection site shows robust axonal expression and limited leaky soma labeling. **G** Native ChR2-tdTomato fluorescence in the motor thalamus (Ventral Anterior Lateral nucleus, VAL) primarily reflects axonal labeling, with minimal leaky somatic expression, contrasting with the extensive somatic labeling observed with the AAV9 intersection strategy.

Manual cell counting revealed robust on-target expression in the right cortex and minimal off-target expression in the left cortex, based on epifluorescence images across the anterior-posterior axis (**Figure 4E**). At the expression peak (+10.4 mm A/P) in the right cortex, 378 Jaws+, 91 ChR2+, and 54 co-expressing cell bodies were identified, corresponding to 14.3% co-expression within the Jaws+ population and 59.3% within the ChR2+ population. In contrast, the left cortex at the same coordinate showed only 26 Jaws+ and 3 ChR2+ cell bodies, along with terminal field labeling in both channels (**Figure 4F**). The expression area in the right cortex extended ∼3 mm along the A/P axis— more focal than the broader spread (∼6 mm) seen with AAV9 (**Figure 2E**). Despite the higher number of Jaws+ somas, labeled Jaws+ axons exiting the region were notably fewer compared to ChR2⁺ axons (**Supplemental Figure 4**). Off-target expression in the thalamus was also substantially reduced, with only two labeled cell bodies observed in the right ventral anterior thalamus, aside from expected axonal labeling (**Figure 4G**). Thus, although AAV8 resulted in moderately lower overall expression efficiency than AAV9, it avoided overexpression-related fluorescence accumulation and significantly reduced off-target and leaky labeling, supporting its suitability for projection-specific targeting.

### Intersectional AAV8-Mediated ChR2 and Jaws Co-Expression in Marmoset Premotor–Parietal Projections

While tests in cross-callosal projections in premotor cortex validated viral efficiency and specificity, we also sought to test this method in a longer-range cortico-cortical projection pathway. We selected a feedback projection from A6DR in premotor cortex to the medial intraparietal area (MIP) in parietal cortex for testing, which is an ipsilateral projection that traverses 13 mm of cortex along the anterior/posterior axis and contains robust feedforward and feedback interconnectivity. Using the same injection strategy as in the cross-callosal experiment above, we injected AAVretro-hSyn-Cre-BFP into the right parietal cortex and a 1:1 mixture of AAV8-CAG-FLEx-ChR2-tdTomato and AAV8-CAG-FLEx-Jaws-GFP into the right premotor cortex (**Figure 5A**).

**Figure 5.**
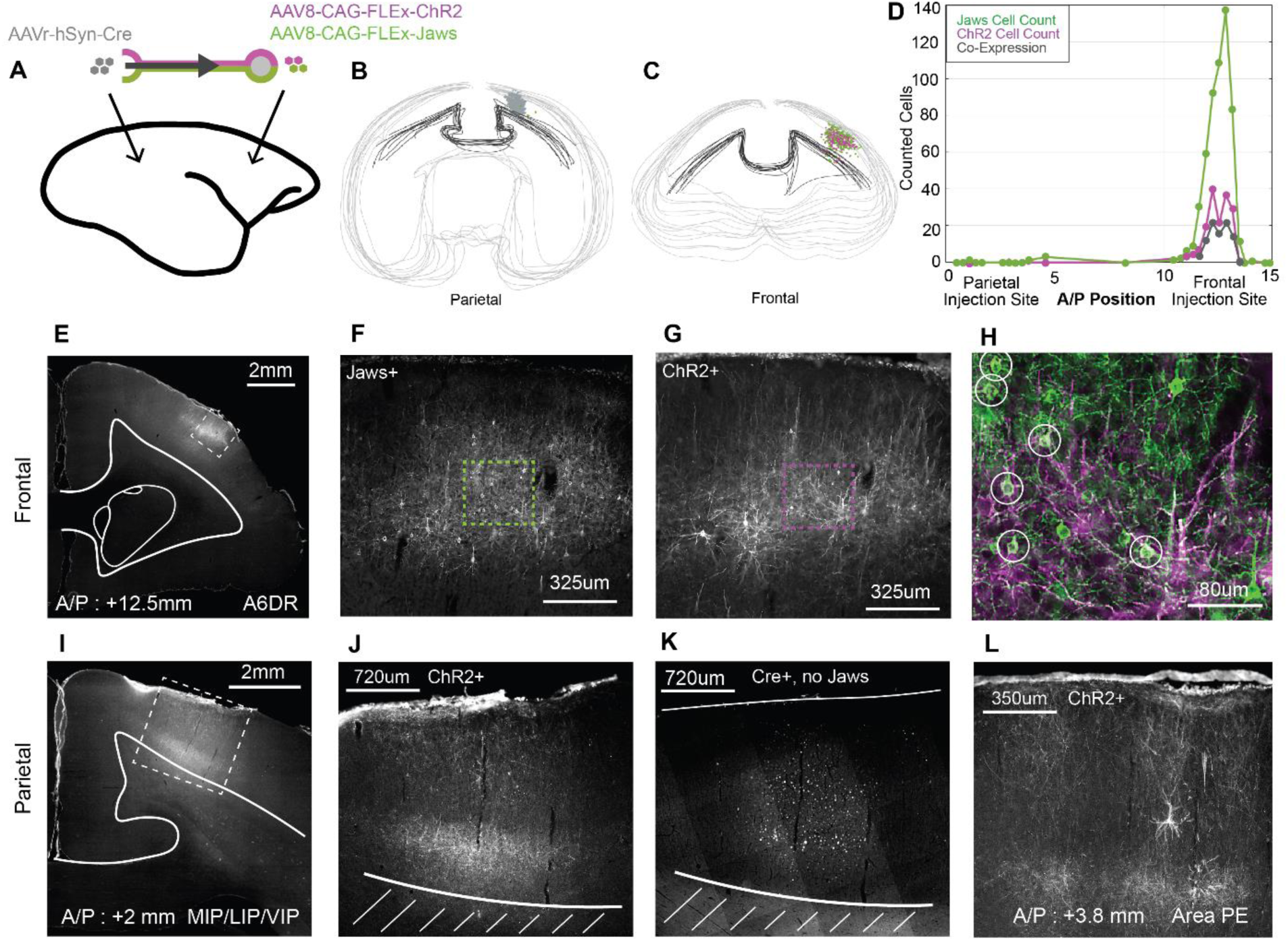
Efficient and Specific Premotor-Parietal Intersectional Expression of ChR2 and Jaws using AAV8. **A** Intersectional viruses were delivered to parietal (AAVretro-Cre) and premotor (AAV8-ChR2 + AAV8-Jaws) cortex in the marmoset. **B–C** Illustration of manual soma counting over coronal sections, with drawn brain perimeter and corpus callosum. Each counted cell is marked as grey (Cre+), green (Jaws+), or magenta (ChR2+). **D** Summary of manual soma quantification across sections, indicating expression of Jaws, ChR2, and co-expression. Each dot represents one counted section. **E** Example section at the center of the premotor injection site showing abundant Jaws+ cell bodies. **F–G** Higher magnification from panel E showing labeled Jaws+ and ChR2+ somata. **H** Two-channel overlay from the region highlighted in F–G, with white circles indicating co-labeled neurons. Jaws+: green; ChR2+: magenta. **I** Example section at the parietal injection center showing dense ChR2+ terminals. **J** Zoom-in from I showing abundant ChR2+ terminals in both deep and superficial layers. Lines mark the cortical surface and white matter boundary. **K** Zoom-in from I showing Cre+ cell nuclei but no detectable Jaws+ axons, based on anti-GFP/anti-BFP immunostaining. **L** Anterior parietal section showing additional ChR2+ terminals, absence of Jaws+ labeling, and a single putative leaky soma.

Consistent with the previous callosal tests, the AAV8 intersection strategy achieved efficient transduction and high specificity for this longer-range frontal-parietal projection. To verify injection sites and assess labeling specificity, we performed manual cell counting to map the 2D positions of labeled Cre+ cells in the parietal cortex and ChR2+/Jaws+ cells in both parietal and premotor cortices, enabling estimation of on-target and off-target labeling (**Figure 5B,C**). Quantification at the premotor site included 15 coronal sections sampled at 0.3 mm intervals, which spanned 5 mm of brain tissue along the A/P axis. A total of 137 Jaws+ cells, 37 ChR2+ cells, and 31 co-expressing cells were counted at the section containing maximum labeling (A/P +12.85 mm, A6DR) (**Figure 5D**). This was equivalent to 22% co-expression in the Jaws+ population and 83.8% co-expression in the ChR2+ population. While cells were counted at all sampled sections, the main area of expression extended from +11.35 mm to +13.45 mm. Qualitatively, labelled cells at the premotor injection site (**Figure 5E**) showed characteristic patterns of healthy expression, with Jaws+ cell bodies appearing more well-defined (**Figure 5F,H**) than those labeled with ChR2. Despite less distinguishable somatic labeling in the ChR2 channel, dense labeling of dendritic processes was still evident (**Figure 5G,H**). An additional feature of expression in the premotor area was the presence of clustered labelled contralateral terminals, suggesting that this population of neurons projecting to the parietal cortex also sends axon collaterals to contralateral premotor areas and more anterior ipsilateral areas (**Supplemental Figure 5**).

We observed high specificity in targeting the projection from premotor to parietal cortex, as indicated by soma labeling restricted to the premotor cortex, with a relative absence of labeled somas in the parietal cortex (**Figure 5D**). To quantify this, 50 μm coronal sections were sampled at 0.5 mm intervals spanning +0.5 to +4.5 mm A/P. At the parietal injection site, very few labeled cells were detected, suggesting minimal off-target expression in the reciprocal projection pathway. Within a 2 mm range centered on the peak of Cre+ expression at A/P +1.8 mm (intraparietal sulcus areas), only one ChR2+ cell and one Jaws+ cell were found. Slightly anterior in parietal Area PE (+3 to +4.5 mm A/P), five Jaws+ and three ChR2+ cells were identified (**Figure 5D**). Imaging at the parietal injection site (**Figure 5I**) revealed dense ChR2+ axon terminal labeling in cortical layers 1 and 5/6 (**Figure 5J**), consistent with the laminar targeting of feedback projections. Notably, Jaws+ axon terminal labeling was absent (**Figure 5K**), suggesting that ChR2 is more efficiently transported into long-range axons than Jaws. This pattern persisted in anterior parietal areas. Although a few cell bodies were detected in these regions (**Figure 5L**), suggesting limited expression leakage in the reverse direction, quantification confirmed that off-target labeling was minimal, accounting for less than 1% of labeled cells (1 of 177 ChR2+ and 5 of 546 Jaws+ cells in the parietal cortex).

To assess the stability and reliability of expression over extended time periods, we repeated the experiment and extended the typical 1.5-month expression window to 8.5 months. This was performed using the same premotor and parietal injection sites, but with ChR2 alone. Histological analysis and cell counting revealed healthy expression in the premotor cortex, with an increased number of labeled ChR2+ cells at the expression peak (113 cells), potentially due to the absence of competition with Jaws. However, this preparation also showed a proportionally higher number of labeled cells in the parietal cortex (peak = 34), indicating increased off-target expression (**Supplemental Figure 6**). Quantification across sections sampled at 0.3 mm intervals showed a mean of 90.37 ChR2+ cells (SEM = 7.84) in the 1.5 mm region surrounding the expression peak in premotor area A6DR (+12.1 to +13.6 mm A/P). In the parietal cortex (+0 to +5 mm A/P), a mean of 8.47 ChR2+ cells (SEM = 2.32) were counted. Overall, the proportion of off-target expression increased to 15% (129 of 754 total ChR2+ cells), compared to just 1% at 1.5 months. These findings suggest that extended expression durations may amplify expression leakage that remains undetectable at earlier time points.

### Excitation and Inhibition of Specific Projection Neurons via Intersectional Optogenetics in the Marmoset

While the intersectional strategy produced clear and specific expression patterns in histological analyses, further validation was necessary to determine whether this level of expression was sufficient to modulate neuronal activity *in vivo*. To move beyond anatomical characterization and assess functional efficacy, we tested the excitatory and inhibitory effects of this intersectional labeling approach through in vivo electrophysiology. Recordings were performed in an anesthetized marmoset with cross-callosal co-expression of ChR2 and Jaws (**Figure 4**). Optical stimulation followed a protocol similar to that used in mice: 5 Hz, 50 ms ChR2 pulses (470 nm, 127.32 mW/mm² at tip) were delivered to the cortical surface, along with simultaneous 1 Hz, 400 ms Jaws pulses (561 nm, 159.15 mW/mm²; **Figure 6A**). To improve the yield of spontaneously active units, we used a combined dexmedetomidine and isoflurane anesthesia protocol (see **Methods**), allowing for reduced isoflurane concentrations. Electrophysiological signals were filtered and spike-sorted to identify units showing optogenetically modulated firing rates.

**Figure 6.**
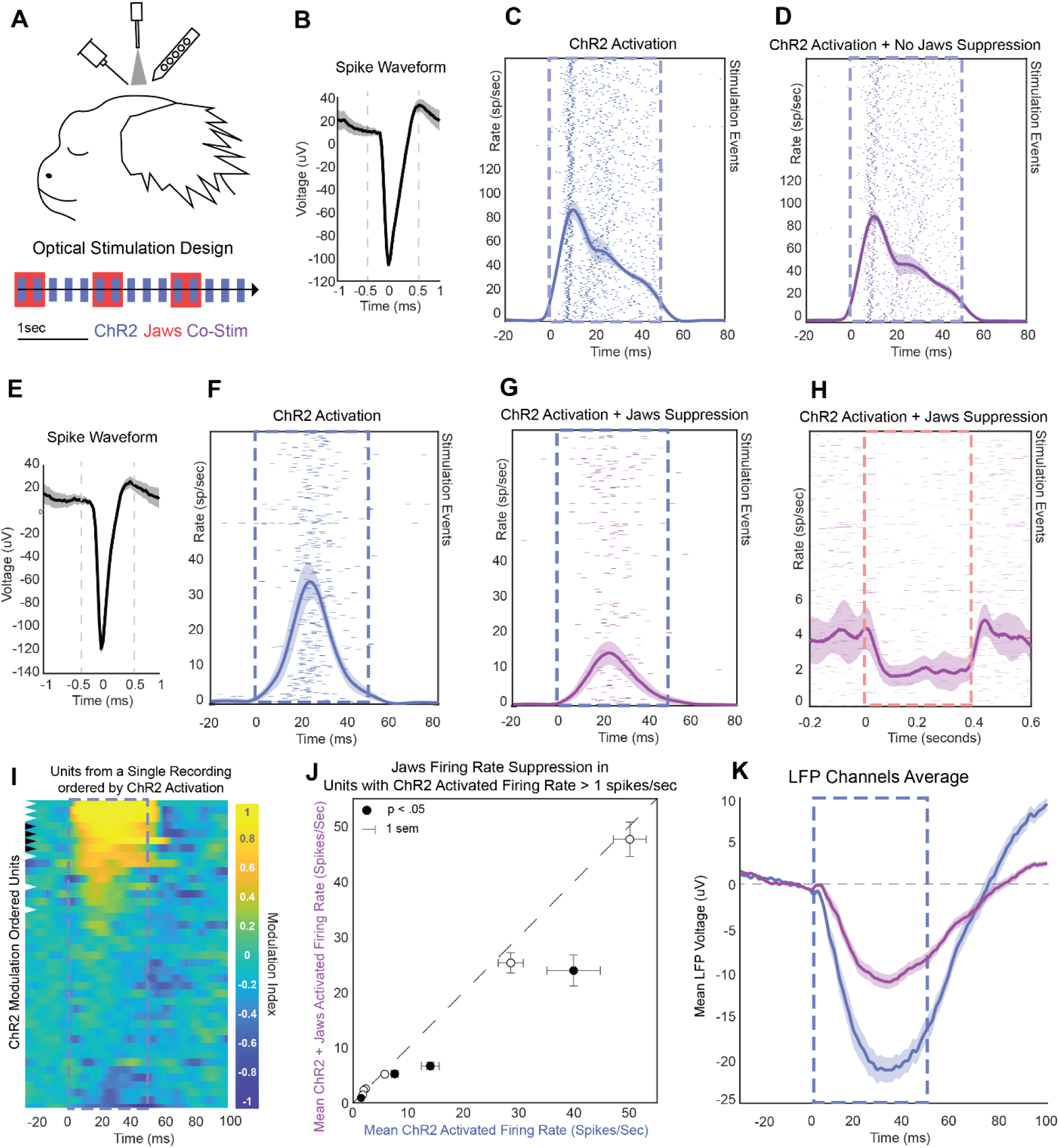
Excitation and Inhibition of Projection Neurons via Intersectional Co-Expression of ChR2 and Jaws. **A** Optogenetic testing was performed in an anesthetized marmoset. Corresponding histological data for this experiment are shown in Figure 5. Optical stimulation included 50 ms pulses of 480 nm light for ChR2 activation (blue, bottom inset), delivered either alone or in combination with 400 ms pulses of 561 nm light for Jaws activation (red), yielding ChR2-only or ChR2+Jaws co-stimulation conditions (purple). **B–D** Spike waveform and peri-stimulus time histogram (PSTH) for a single unit showing robust firing in response to ChR2 stimulation and minimal suppression during co-stimulation with Jaws. Shaded areas represent ±2 SEM. **E–G** A second example unit in the same format as B–D, showing marked suppression of ChR2-evoked firing during concurrent Jaws activation. **H** Time-locking this unit’s activity to the 400 ms Jaws pulses during co-stimulation reveals significant firing suppression relative to baseline. **I** Heatmap of ChR2 modulation effects across all units (yellow = excitation, blue = suppression), ordered by degree of ChR2-evoked modulation. Black and white arrowheads indicate the example units shown in B–G. **J** Scatterplot comparing the mean firing rate during ChR2-alone vs. ChR2+Jaws co-stimulation for units that exhibited >1 Hz ChR2-evoked increases over baseline. Black circles indicate units with significantly reduced firing during co-stimulation. **K** Mean population local field potential (LFP) traces show that Jaws co-stimulation reduced the ChR2-evoked LFP deflection by 62% at peak amplitude, indicating effective population-level suppression.

A subset of neurons, presumed to be neurons projecting to the contralateral premotor cortex, exhibited clear responses to optogenetic stimulation. The extent of suppression by Jaws co-stimulation varied across units, ranging from no suppression to nearly 50%. One example unit showed short-latency, laser-triggered activation consistent with direct ChR2-driven excitation, but its firing remained unaffected during concurrent Jaws stimulation (**Figure 6B–D**). By contrast, another unit displayed ∼50% suppression of ChR2-evoked firing during simultaneous Jaws stimulation (**Figure 6E–G**). Time-locking this unit’s activity to the 400 ms Jaws stimulation pulses alone also revealed clear suppression of the ChR2 stimulation-enhanced firing rate (**Figure 6H**). Out of 67 units recorded in a single session, 10 showed significant modulation of ChR2-evoked firing above 1 Hz. Population-level responses are summarized using a modulation score as described previously (**Figure 6I**). Analysis by cortical depth revealed modulated units at the deepest extent of the electrode array, confirming successful activation of projection neurons located in deep layers despite surface laser stimulation (1.1 mm; **Supplemental Figure 7**). Among the 10 ChR2-responsive units, four were significantly suppressed by concurrent Jaws stimulation, while none showed significant excitation (p < 0.05, two-sample, two-tailed t-tests comparing firing rates during ChR2 vs. ChR2+Jaws within a 5– 45 ms window post-ChR2 onset; **Figure 6J**). These suppressive effects were also reflected in the local field potential (LFP). A comparison of mean LFP traces showed a 62% reduction in the ChR2-evoked peak voltage deflection post-laser onset during concurrent Jaws activation (**Figure 6K**). Although the number of modulated neurons was small, as expected given the low proportion of callosal projection neurons in the population, these results confirm effective ChR2-mediated excitation and partial suppression by Jaws in targeted projection pathways.

## Discussion

Successful application of optogenetic techniques in non-human primates remains a major challenge, particularly with respect to achieving selective expression in specific cell types or projection pathways. In this study, we optimized viral delivery strategies to achieve highly specific optogenetic modulation restricted to defined cortico-cortical projection pathway in the common marmoset. We first validated our optogenetic preparation by replicating a well-established inhibitory approach previously used in macaques to induce behavioral deficits^20,21^. Using the mDlx promoter to drive ChR2 expression in inhibitory interneurons, we confirmed that surface cortical stimulation suppressed local neural activity. Spike waveform analyses further indicated activation of deep-layer inhibitory interneurons. Next, we developed and optimized an intersectional strategy for projection-specific targeting in marmoset cortex, using both mouse and marmoset models to identify optimal AAV serotypes and titers. This approach enabled target-specific expression in both shorter-range callosal projections between left and right premotor cortices and longer-range frontal-to-parietal projections within a hemisphere. Moreover, we successfully co-expressed two opsins, ChR2 for excitation and Jaws for inhibition, within the same population of projection neurons. Following optimization, this strategy yielded stable, healthy expression in marmoset cortical tissue. Electrophysiological recordings confirmed that the intersectional approach produced sufficient expression to drive optogenetic modulation of neural activity *in vivo*.

Optimizing viral expression methods to enhance targeting specificity without compromising efficiency is essential for functional neuroscience experiments. Without conditional expression strategies like those described here, low specificity can result in modulation of heterogeneous neural populations, comprising mixed cell types and projection targets, which complicates interpretation. For example, we found that expressing ChR2 using a highly efficient virus like AAV9 combined with a strong pan-neuronal promoter resulted in widespread labeling of local, retrograde, and anterograde projections in marmoset, consistent with previous rodent studies^27,29^. Such broad labeling leads to ChR2 expression in diverse neuronal populations, making it impossible under standard optical stimulation to selectively activate only the projection neurons of interest without concurrently stimulating interneurons, inducing antidromic spikes, and triggering widespread, mixed network effects. Classical approaches, including microelectrode stimulation, pharmacological modulation, surgical lesions, and cortical cooling, suffer from similar limitations. Recent innovations, such as soma-targeting sequences, can reduce axonal activation and antidromic firing in distantly labeled neurons^53,54^. Additionally, 3D optical targeting strategies using multiphoton excitation offer enhanced spatial control^55^. However, these methods remain either technically demanding or incomplete solutions. Given the current lack of transgenic Cre-driver lines in non-human primates, two-virus intersectional strategies represent one of the most effective available tools for achieving projection-specific targeting, enabling the functional dissection of defined neural circuits in the primate brain.

To optimize intersectional viral expression, we first conducted testing in mice before proceeding to marmosets. Each viral batch was initially evaluated in the mouse, allowing for preliminary screening of injection parameters and the identification of key methodological adjustments. Notably, we found that dilution of the locally injected AAV8 virus was necessary to prevent overexpression. This dilution likely reduced the concentration of residual cytotoxic components from the viral preparation process and/or limited the number of viral particles entering each cell, thereby preventing metabolic stress at high concentrations^56^. This step proved critical, both for refining our procedures and for reducing the number of marmosets required for the study. Moreover, the mouse-based screening allowed us to compare multiple AAV serotypes and ultimately identify AAV8 as the most suitable candidate due to its lower retrograde transport affinity, which minimized off-target expression in the reciprocal projection pathway.

Translating intersectional methods to the marmoset model required further adjustments to our injection strategy to accommodate the species’ larger brain size and anatomical variability. This variability, well-documented across individuals, poses a particular challenge for stereotaxic targeting in the absence of MRI guidance^57^. To increase the likelihood of targeting the intended regions, we distributed injections across multiple sites within each craniotomy. Additionally, higher injection volumes and multiple depth penetrations were used to minimize issues such as backflow and uneven viral distribution. An interesting source of variability observed in AAV8 experiments was the differing number of Jaws-expressing versus ChR2-expressing cells. One possible explanation is viral competition, in which infection by one vector may reduce the efficiency of co-infection by the second, thereby decreasing the rate of co-expression. Alternatively, the potassium channel trafficking motifs included in the engineered Jaws construct may preferentially localize expression near the soma^52,58,59^, potentially biasing detection in cell counts.

While AAV9 demonstrated high transduction efficiency in our initial intersectional tests, consistent with previous reports^38^, it also produced leaky labeling, likely due to its known retrograde transport affinity^29,45^. A prominent finding was the abundance of off-target soma labeling in the left cortex and in the motor thalamus, relative to the targeted right cortex. Although anterograde trans-synaptic transport has been reported for AAV1 and AAV9^28^, to our knowledge, this property has not yet been documented for AAVretro. Alternatively, the thalamic labeling may result from retrograde transduction by both AAVretro and AAV9 in thalamic neurons that project to the right premotor cortex and also send decussating axons to the left premotor cortex. Such decussating thalamo-cortical projections have been described in the rat^48^, hedgehog^60^, and macaque^46^. Importantly, our data suggest that AAV9 can still be used effectively without overexpression when paired with cell-type-specific promoters. For example, when combined with the inhibitory neuron-specific mDlx promoter, AAV9 yielded healthy, specific labeling that remained stable for at least 10 months. These findings demonstrate that AAV9 remains a useful tool when paired with appropriate promoter systems that constrain moderate expression in specific neuronal populations.

The application of mDlx-ChR2 in the marmoset cortex confirmed its utility for silencing activity within a local cortical area. Previous studies in macaques have used mDlx-driven optogenetics to achieve cortical inhibition and demonstrated robust electrophysiological inactivation in primary visual cortex (V1), with modulation profiles closely resembling those observed in our current results^21^. Our findings contribute two important advances to ongoing research on mDlx-mediated optogenetics. First, we employed a noninvasive method of surface illumination through a silastic-protected cranial window. Combined with single-unit spike waveform analysis, this approach enabled the detection of inhibitory interneuron activation at depths of up to 1.1 mm from the cortical surface. This demonstrates the feasibility of noninvasive optogenetic activation of deep-layer neurons, which is particularly valuable for experiments not involving invasive optical delivery. Second, our study extends the use of mDlx-ChR2 to the premotor cortex, a region with fundamentally different laminar architecture from V1. In contrast to the laminar-specific expression pattern previously observed in macaque V1, including a clear absence in Layer IV, our results showed no discernible laminar bias in premotor cortex. This difference likely reflects both regional and species-specific variability in interneuron distribution and viral expression. Further comparative studies will be important to clarify how these anatomical factors influence mDlx-based optogenetic labeling across brain regions and species.

The direct excitatory and inhibitory optogenetic methods explored in this study achieve high labeling specificity through a viral intersection strategy; however, this approach has certain limitations. When targeting long-range projections, stimulation at axon terminals can help avoid off-target effects associated with optical stimulation at the soma site. For example, a study of feedback projections from the frontal eye field in macaques successfully achieved such specificity by stimulating projection terminals with the inhibitory opsin eNpHR3.0^32^. This approach, however, requires robust expression throughout the entire projection pathway, an essential criterion that was clearly met by ChR2 in our experiments, but not consistently by Jaws. In the premotor-parietal projection, we observed no Jaws-positive labeling in the parietal cortex. Although Jaws-positive terminals were present in the premotor cross-callosal projection, their expression appeared less robust than that of ChR2. These observations suggest that Jaws is less efficiently transported to, or incorporated into, distal axonal membranes. A recent study employing Jaws-mediated terminal inhibition in a macaque V4-to-V1 feedback projection demonstrated functional inhibition via electrophysiology^31^, but did not provide histological characterization of Jaws expression at terminals, leaving unresolved the question of its transport and localization efficiency in long-range projections. It remains possible that inherent features of the Jaws construct limit its trafficking and efficacy in distal axons, warranting pathway-specific validation for terminal inhibition applications. Alternative tools, such as opto-GPCR-based inhibitory systems^61^ that suppress synaptic transmission, may offer more reliable options for long-range projection targeting and may be considered to overcome this limitation. In addition, the intersectional viral labeling methods could also be combined with chemogenetic constructs to enable selective modulation of projection-defined neuronal populations with systemically delivered drugs^33,62^.

While the co-expression and two-color optogenetic approach used in this study enabled simultaneous excitatory and inhibitory modulation in the anesthetized marmoset, there is room for improvement in the co-injection strategy. Jaws has previously been applied successfully in primate studies for site-specific inhibition, including in the superior colliculus^63^ and area MT^22^. Similarly, ChR2 has seen broad application in primate optogenetics, from evoking forelimb movements via M1 stimulation^64,65^ to modulating reward signaling in an amygdala–substantia nigra circuit^66^. In our preliminary mouse experiments, Jaws effectively suppressed ChR2-driven activity, with some units showing near-complete inhibition. In contrast, the suppression observed in the marmoset was more subtle. One potential explanation is competition between the ChR2 and Jaws vectors for expression within the same neurons. As shown in our histological data and electrophysiological recordings, co-expression was not always well matched across cells, suggesting that complete bidirectional optogenetic control is not reliably achieved with this dual-virus method at the single neuron level. Given the likelihood of viral competition, a promising alternative is the use of a single opsin construct that combines excitatory and inhibitory components into a single molecule. One such tool, BiPOLES^67^, enables bidirectional modulation via dual-wavelength stimulation and may offer a more reliable and efficient solution for future studies.

The intersectional optogenetic methods described here expand the growing toolkit available for primate optogenetics. To date, much of the field has focused on well-characterized or predominantly unidirectional circuits, such as those in the frontal eye fields^25^, superior colliculus^63,68^, and motor cortex^64^. By employing a viral intersection strategy, our approach enables the selective targeting of individual components within complex, integrative cortical circuits, such as those found in associative areas, with minimal off-target expression. These findings also highlight the utility of multi-color optogenetics, particularly when combined with cell type–specific tools. Used in tandem, these approaches allow researchers to replicate the functional outcomes of classical perturbation techniques, such as cortical cooling or muscimol injection, but with the added temporal precision and specificity of optogenetics. Moreover, selective projection inhibition from a defined source area offers a powerful contrast to local inactivation, helping to disentangle circuit-level contributions to behavior and physiology. Importantly, this work also validates a minimally invasive method for delivering light through a cortical window in the primate brain, further broadening the methodological options for chronic or awake preparations. Marmosets possess highly developed visual systems and exhibit cooperative social behaviors, making them particularly well-suited for functional and behavioral studies in primates^69–73^. Our results demonstrate the feasibility of selectively targeting projection pathways between reciprocally connected cortical areas in primates and provide a robust framework for probing their causal role in brain function.

## METHODS AND MATERIALS

### Animal Use

All experimental protocols were approved by the University of Rochester Institutional Animal Care and Use Committee and were conducted in compliance with the National Institutes of Health guidelines for animal research.

Twenty-six wild type C57BL/6J mice were used for this study. Twenty (12 female, 8 male) were used for virus injection and histology alone. Three (3 male) were used for virus injection, anaesthetized terminal electrophysiology recordings, and histology. Three (3 female) were used for virus injection, awake head-fixed electrophysiology on a run wheel, and histology. Subjects were group housed at the University of Rochester Medical Center with a circadian cycle of 12-hour light and 12-hour dark. All subjects had full access to food and water at all times.

Five adult common marmosets (Callithrix jacchus) were used for this study. Marmoset S (male) and Marmoset X (female) were used for virus injection and histology alone. Marmoset Y (female) and Marmoset K (female) were used for virus injection, a terminal anaesthetized electrophysiology recording, and histology. Marmoset A (male) was used for virus injection, awake, head-fixed electrophysiology recordings, and histology. A summary of these experiments is provided in Table 1. Subjects were group housed at the University of Rochester with a circadian cycle of 12-hour light and 12-hour dark. All subjects had full access to food and water at all times.

**Table 1.**
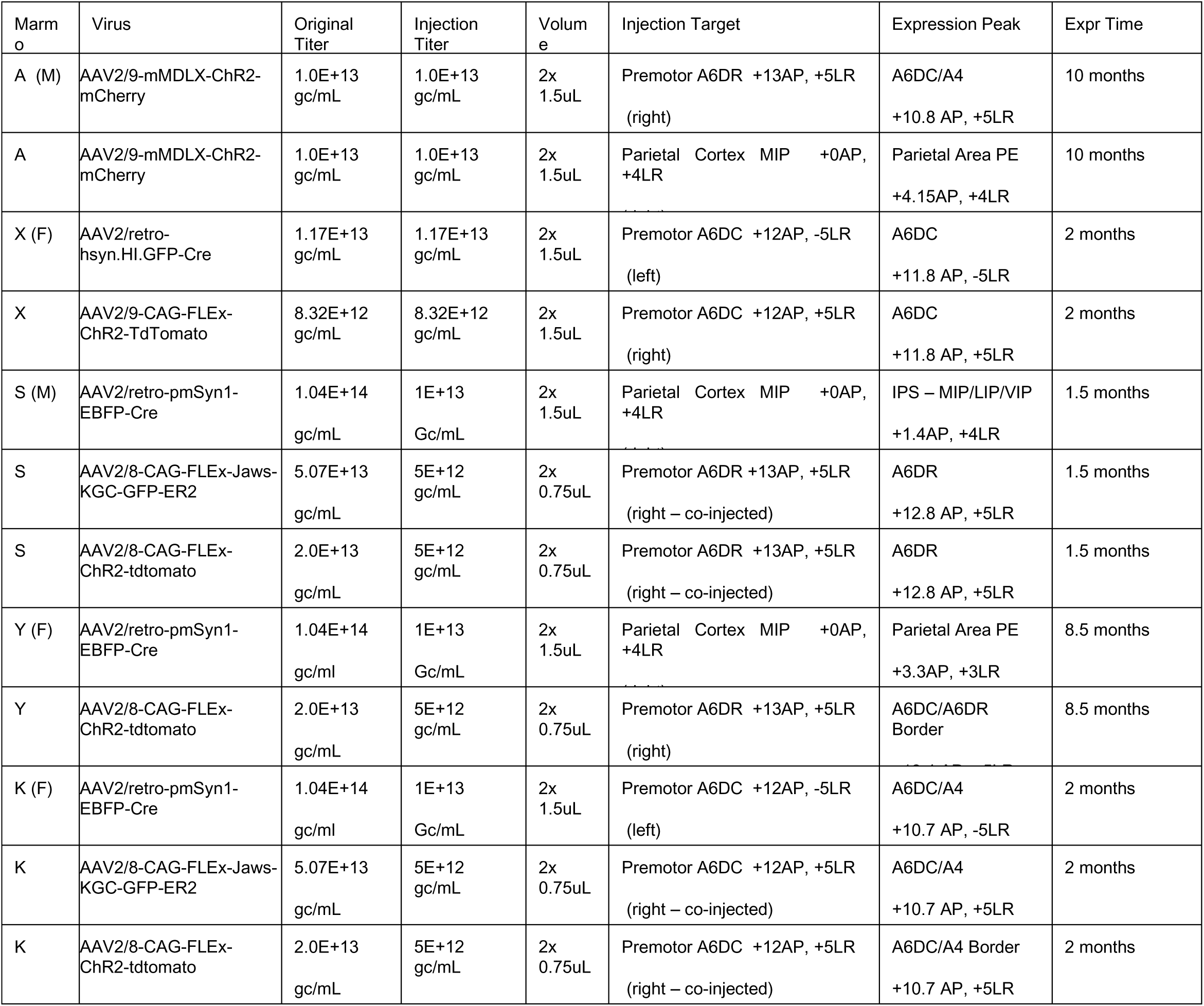
Marmoset Injection Procedures.

**Table 2.** Viral Materials.

| Virus | Plasmid Manufacture | Virus Manufacture |
| --- | --- | --- |
| AAVretro-hsyn-cre-GFP | Addgene | Addgene #105540-AAVrg |
| AAV2/retro-pmSyn1-EBFP-Cre | Addgene #51507 | Boston Children's Hospital (BCH) Viral Core |
| AAV2/9-CAG-ChR2-tdtomato | BCH Viral Core | BCH Viral Core |
| AAV2/8-CAG-ChR2-tdtomato | Addgene #18917 | BCH Viral Core |
| AAV2/8-CAG-FLEX-Jaws-KGC-GFP-ER2 | Addgene #84445 | BCH Viral Core |
| AAV2/9-MDLX-ChR2 | Addgene | Addgene #83898-AAV9 |

### Marmoset Injection Surgery

Prior to surgery, a final virus solution was prepared by mixing sterile saline with undiluted virus aliquots to achieve titers described in Table 1. Note that for co-injected virus (e.g. Jaws and ChR2), individual virus solutions were diluted to 1E+13 genomic copies per mL (gc/mL) and then mixed 1:1 in the final injected solution.

Prior to viral injection, marmoset research subjects were anaesthetized with intramuscular injection of Ketamine and Midazolam followed by isoflurane delivery using a full-face mask prior to endotracheal intubation. After intubation, anesthesia was maintained using 1.5-3% isoflurane while monitoring blood oxygenation, heart rate, rectal temperature, and respiration. The head was fixed to a Kopf stereotaxic frame using ear bars and a Kopf Model 916 Guinea Pig adaptor nose and tooth bar assembly. The head was positioned on the tooth bar to distribute pressure over the left and right sides of the dental arch. The head was rotated about the ear bars such that the plane formed by the lower orbits and the ear bars was parallel to the base of the stereotaxic frame. A stereotaxic arm was fixed to the frame and calibrated to align the x and y planes with respect to the ear bars to mark stereotaxic coordinates. An incision was made in the scalp and tissue was cleared at the skull surface. Injection target locations were marked on the skull (Premotor-Premotor: 10 mm anterior to intra-aural (IA) line, 4 mm left and right of midline; Premotor-Parietal: 10 mm and 0 mm anterior to IA, 4mm right of midline). 3 mm moat craniotomies were drilled at each target site. The skull chip was removed, and craniotomy edges were smoothed with a coarse Dremel bit.

Two injection site targets were identified within each craniotomy based on the pattern of vasculature. A small slit was made in the dura at each target. A Nanoject III (Drummond Scientific) was set to inject 3 quantities of 500 nL of viral solution at each dural slit at cortical depths of 1.5 mm, 1 mm, and .5 mm. The injection rate was set to 2nL/second. Two minutes were allowed to pass before the beginning of each injection and after the end of each depth injection. After injection, craniotomies were sealed with a thin layer of bone wax and the skin was sutured following the administration of slow-release analgesic for post operative recovery. A minimum expression period of 6 weeks elapsed prior to sacrifice for histology or electrophysiology experiments. Further experimental details for individual animals are documented in Table 1.

### Marmoset Head Implant and Injection Surgery

Marmoset A was surgically implanted with a head cap and recoding chambers for awake, head-fixed optogenetic stimulation and electrophysiology. Two months prior to surgery, Marmoset A was trained to sit in a small primate chair following methods previously described^74,75^. Under sterile conditions, Marmoset A was implanted with an acrylic head cap with titanium posts to stabilize the head using methods described in detail previously^74^. During the implant surgery, recording chambers were placed over premotor area A6DR and parietal area MIP based on stereotaxic coordinates^76^. Custom 3D printed (Proto Labs) recording chambers were adhered to the skull using C&B-Metabond (Parkell, Inc.). The skull exposed within each recording chamber was covered by a thin layer of C&B-Metabond.

After recovery from head implant surgery, Marmoset A was trained to acclimate to head-restraint while sitting comfortably in a custom designed primate chair. Over several months, he was trained to perform several basic tasks including central fixation^77^, saccade tasks towards peripherally detected stimuli^74^, and free viewing of natural images. After preliminary training, a second surgery was performed to create craniotomies and perform virus injections in both recording chambers according to the procedures described above. Isoflurane anesthesia was delivered with a custom 3D printed (Proto Labs) marmoset anesthesia mask rather than using endotracheal intubation. This mask was designed to fit onto the Kopf bite bar of our stereotaxic frame and formed a seal with the face using a custom cut sheet of clear, food grade silicone rubber.

Craniotomies were sealed with a thick layer of silastic gel (Kwik-Sil; World Precision Instruments) to protect the brain from infection and reduce granulation growth^78^. Post-operative recovery from craniotomies required removal of silastic and flushing the craniotomy with sterile saline to remove coagulated blood until the tissue stabilized. After stabilization, the dura at each craniotomy site was excised under sterile conditions. A thin layer (<1 mm) of Silastic was then applied to seal the open craniotomy, which was thin enough to enable passage of electrodes for recordings. The silastic remained in place for the duration of the study. When applied correctly, the silastic has been observed to limit the growth of granulation tissue, prevent infection^78^, and can also be penetrated by linear array silicon probes with minimal risk of damage to the probes. Further, penetration sites are naturally sealed by the Silastic after removal of the electrodes.

### Awake and Anaesthetized Marmoset Electrophysiology and Optogenetic Stimulation

A 64-channel silicone array (NeuroNexus, Inc.) was used to record neural signals during optogenetic stimulation while Marmoset A free-viewed natural images under head fixation^79^. Sub-millimeter depth control of the array was achieved using a custom-built X-Y stage. The stage mounted onto the recording chamber and carried a light-weight screw microdrive (Crist Instrument, 3-NRMD). The 64-channel array consisted of 2 probe shanks separated by 200 microns. Each shank consisted of 32 channels with 35 microns spacing between contacts. All arrays were 50 microns thick and had sharpened tips. For the best recording quality, arrays were electro-plated with PEDOT, a method that has been shown to increase signal/noise ratios^80,81^.

To deliver light pulses for optogenetic stimulation, a guide tube was built into the X-Y stage oriented to direct a fiber optic cannula to a point on the silastic barrier immediately adjacent to the penetration point of the electrode array. An uncleaved fiber optic cannula (ThorLabs – 400 um core, .39 NA) was cut using an optical scribe to a length whereby the end of the probe touched the silastic without applying pressure when fully inserted into the guide tube.

In preparation for awake electrophysiology recording, Marmoset A was seated in a primate chair and head fixed. The head stage was then mounted onto the premotor chamber, the fiber optic cannula was inserted into the guide tube, and the electrodes were lowered into cortex while viewing electrophysiological signals (Open Ephys GUI). Initial array insertion into cortex followed a slow “two turns in, one turn out” procedure (each turn of the microdrive is equivalent to 250uM), which prolonged the insertion process to occur over a span of ∼30 minutes. This procedure prevented silastic and cortical dimpling-related unit activity suppression and movement of units along the array during recording. All recordings were amplified and digitized at 30KHz using an Intan recording headstage. When units were apparent across all channels in the array (typically at a depth of ∼1 mm-1.5 mm), optogenetic ChR2 stimulation experiments began. During recordings, Marmoset A free-viewed natural images while receiving periodic Ensure from a tube placed in the mouth and an automated syringe pump. Data were collected with this animal over 10 recording sessions spanning a month.

Optogenetic stimulation in awake and anaesthetized recording preparations in the marmoset was conducted using two general protocols for ChR2 stimulation and Jaws stimulation. In all cases, stimulation was delivered from a 400 um core diameter, 0.39 NA fiberoptic cannula positioned closely over the cortical recording site, usually with a slight angle oriented towards the electrode array at depth. Laser pulses were controlled with a custom-built Arduino device capable of sending four independent TTL pulse trains from BNC connectors. In cases of simultaneous Jaws and ChR2 stimulation, two channels were dedicated to controlling laser pulses and the remaining two channels sent encoded laser metadata to the recording system. In the case of ChR2 stimulation, 40-50 ms blue light pulses were delivered with a frequency of 5 Hz. For each pulse, a jitter variable was set to randomly choose a value from 0-75 ms and add or subtract this value from the otherwise periodic pulse times. For Marmoset A (**Figure 1**) and K (**Figure 6**), blue light was delivered from a 470 nm LED with a measured output of 16 mW (470 nm, 127.3 mW/mm^2^ at tip). Additional output powers were tested for mDlx-ChR2 titration: 8 mW (63.6 mW/mm^2^) and 4 mW (31.8 mW/mm^2^). In mouse experiments, blue light was delivered from a 473 nM solid state diode-pumped crystal laser (CrystaLaser) with output powers of 5 mW (39.79 mW/mm^2^), 10 mW (79.57 mW/mm^2^), 15 mW (119.36 mW/mm^2^), or 20 mW (159.15 mW/mm^2^). In experiments using an awake recording preparation (Marmoset A - **Figure 1**), light travelled through at least 1 mm of clear silastic material, thus attenuating the light power by more than 70% of the values measured at the fiberoptic tip by geometric light falloff alone before hitting the brain surface^82,83^. In anaesthetized experiments (Marmoset K – **Figure 6**), the fiberoptic was usually 0.5 mm from the brain surface, which would attenuate intensity values by more than 50% at the brain surface^82^. In the case of Jaws stimulation, 400 ms light pulses were delivered with a frequency of 1Hz. Similarly, a jitter value ranging from 0-200 ms was added or subtracted from otherwise rhythmic pulse times to avoid any intrinsic entrainment of neural activity. For Marmoset K and mouse experiments, 561 nM light was delivered from a solid-state diode-pumped crystal laser (CrystaLaser) with output power ranging from 5 mW to 20 mW. To test the suppression of ChR2-evoked neural activity by Jaws, the above protocols were executed simultaneously from two separately controlled light sources and cannulas. In this configuration, neural activity induced by ChR2 events within the Jaws stimulation window was compared to ChR2 events outside of simultaneous Jaws stimulation.

Anaesthetized marmoset electrophysiological recordings were conducted with Marmoset K (**Figure 6**) six weeks after viral injection surgery. Prior to surgery, a custom electrode array armature was built out of acrylic, metal, and epoxy, which could be attached to a stereotaxic arm. This device was used for mouse recordings as well. Marmoset K was anaesthetized using isoflurane delivered through a custom marmoset anesthesia mask described above. Marmoset K was stabilized in a stereotactic frame using ear bars and a bite bar. An incision in the scalp was made and the premotor ChR2+Jaws injection craniotomy created for viral injection was re-opened. The edges of the craniotomy were expanded using a dental rotary tool, which aided in granulation tissue removal. Once the craniotomy was clear and dura was exposed, the dura was excised using a hooked syringe tip. A fiber optic cannula held by a stereotaxic arm was lowered and inserted into the small pool of saline maintained in the craniotomy. The electrode array was lowered into the brain and electrophysiological recordings with ChR2 optogenetic stimulation were conducted as described above.

To increase yields of recorded units when using isoflurane alone in preliminary anaesthetized experiments, an anesthetic protocol was developed to include, in addition to isoflurane, a dose of buprenorphine HCl (.01-.02 mg/kg) and a loading dose of dexmedetomidine (2.5 ug) followed by a continuous drip of dexmedetomidine (5-40 ug/kg/h)^84^. This allowed an overall reduction in concomitant isoflurane delivery, resulting in increased background cortical activity and general excitability. In this case, the animal was intubated to facilitate manual pumping of breaths in the case of severe respiratory depression. Anesthesia for Marmoset K was titrated using this protocol and a stable plane was achieved using .25% isoflurane and 40 ug/kg/hr of dexmedetomidine. After the animal was stabilized, ChR2 and Jaws optogenetic stimulation and electrophysiological recordings were conducted as described above. After anaesthetized recording procedures, subjects were immediately euthanized and perfused according to procedures described below.

### Mouse Procedures

Injection surgeries and optogenetic electrophysiology recordings in the mouse were conducted to perform initial tests of viral expression, dilution titrations of viral solutions, and to, in general, pioneer methods to be translated to the marmoset. Injection surgeries were performed as above, but with significantly lower volumes (1x 200-400 nL per anatomical area injected at one site over 3 cortical depths) and largely targeting the bilateral M1/M2 projection (Bregma +1.5 mm AP, +/- 1.5 mm LR), as above in the marmoset targeting of bilateral premotor cortex, areas A6 dorsal rostral and A6 dorsal caudal (A6DR/A6DC). Tests of potential spontaneous recombination in viral vectors were conducted in M1/M2 or in posterior vision areas (Bregma -2.5 mm AP, +/-2 mm LR). In these experiments, AAV2/retro-pmSyn1-EBFP-Cre was injected into left M1/M2 and AAV2/8-CAG-FLEx-Jaws-KGC-GFP-ER2, AAV2/8-CAG-FLEx-ChR2-tdtomato, or a 1:1 combination of the two were injected into right M1/M2. Following initial histological injection tests of these vectors, 1:1, 1:2, 1:5, and 1:10 dilutions of these viral solutions were tested. For animals intended for awake electrophysiology procedures, injection procedures followed the implantation of a 3D-printed headframe to the skull using previously described procedures^85^. Following implantation and injection surgery, craniotomies were sealed with Silastic material as described above. Following an expression time of 1.5 weeks, animals were sacrificed for histology or used for electrophysiology experiments.

Mice that participated in awake electrophysiology experiments were acclimated to a head fixed, freely-rotating run wheel apparatus^85^ during the 1.5 week expression time. After this period and prior to recording, mice were first anaesthetized using 1-2% isoflurane to open craniotomies and perform granulation tissue removal if necessary. They were immediately head fixed onto the run wheel apparatus, where they recovered from anesthesia. A stereotaxic arm-held electrode array, as described above, was lowered into cortex while recording signals using the Intan RHD2000 software. Fiber optic cannulae were placed in position directly above cortex for ChR2 and Jaws stimulation during recordings. The electrode and cannulae were moved between ChR2/Jaws and Cre injection craniotomies to test signals at axon terminal sites and the effects of terminal stimulation. At the end of a recording session, craniotomies were re-sealed with silastic. No more than 3 recording sessions were performed per animal. In the case of anaesthetized electrophysiology experiments, scalp incisions were sutured post injection surgery. Animals were anaesthetized using 1-2% isoflurane and secured into a stereotaxic device using ear bars and a bite bar. An incision was made in the scalp and tissue was cleared from injection craniotomy sites. Electrode arrays and cannulae were put in position and isoflurane was lowered while measuring breaths per minute and monitoring for signs of arousal. Once a stable anesthetic plane was found using the lowest concentration of isoflurane (.5%-1%), laser stimulation protocols were tested while recording electrophysiology signals. For these experiments, .5 Hz 1-second duration Jaws stimulation pulse trains were delivered with concomitant 5 Hz 50 ms duration ChR2 pulses.

### Histology

At the termination of electrophysiology experiments or after sufficient virus expression time transpired in the mouse and marmoset, animals were transcardially perfused with 1X PBS or saline solution followed by 4% paraformaldehyde in 1X PBS or 10% formalin solution. Mouse tissue was post-fixed overnight and coronally sliced in 50um-80um sections on a vibratome prior to mounting and cover-slipping sections as necessary. Marmoset tissue was post-fixed for two days and incubated in sucrose solution until neutrally buoyant in 30% sucrose solution. Frozen coronal slices were cut to a thickness of 50um-80um on a microtome and mounted and cover-slipped as necessary or stored in 1x PBS with .05% azide. All sections were imaged using a confocal microscope (Olympus FV1000) or an automated epifluorescence slide scanner (Olympus VS110). Some fluorescent reporter signals required amplification using immunohistochemistry staining after remaining in storage for 3 months or more. Anti-RFP, anti-GFP, and anti-Cre antibodies were utilized in this study, generally in a 1:1000 concentration with 1-3 marmoset sections in a well and followed by fluorescent secondary antibodies with a 1:100 concentration. Image processing was performed using custom Matlab (2022b, Mathworks) scripts, FIJI 1.54h^86^, QuPath 5.1^87^, OlyVIA 2.9 (Olympus Scientific Solutions), and Photoshop (Adobe).

### Data Analysis

#### Image Processing

In addition to image visualization optimization, viral reporter expression levels were quantified using manual cell counting and automated tissue classification. Manual cell counting was performed on an epifluorescence slide scanner (Olympus VS110) images of 50um-thick marmoset tissue sections. Slide scanner images were used due to their high throughput efficiency. Custom Matlab scripts were used to first separate color channels of interest, apply a square root transformation to each image matrix, and perform levels corrections. Images were imported into Image J and manually labelled using the Cell Counter tool. Opsin positive cell bodies were identified by the characteristic hollow spheroid shape of cell membrane labelling. Slightly out of focus cell bodies were included in this counting procedure. For co-expression counting, two-channel images were imported into ImageJ and any evidence of overlapping spheroid-shaped labelling was counted as a positively co-expressing cell body regardless of differences in labelling intensity. For the AAV9 intersection experiment involving Marmoset A, the intensity and density of neuropil labeling made it impossible to perform accurate cell body counting. Therefore, QuPath was used to perform automated tissue classification using an in-built artificial network and a training procedure that provided the network with training images of examples of no tissue, un-labelled tissue, labelled tissue with healthy expression, and labelled tissue with unhealthy over-expression. One-channel images were scaled to 775 px/mm, baseline subtracted, square root transformed, and intensity corrected to bring the scope of expression intensity within display range. Pre-processing in QuPath consisted of .5px gaussian smoothing to emphasize sub-cellular features and 4x gaussian smoothing to emphasize broader features. Principal components analysis feature normalization was also performed. Training images were submitted to a 5-layer artificial neural net with a sigmoid activation function. For training image selection, the distinction between healthy and unhealthy labelled tissue was the clear and distributed presence of fluorescent puncta that likely reflect fluorescent protein aggregates and/or degenerating neural processes. Labelled healthy tissue was defined by spherically labelled cell bodies without puncta and included terminal labelling, such as labelling found in basal ganglia and claustrum. This classification procedure was summarized by dividing the area classified as healthy or unhealthy expression by the sum of areas classified as healthy, unhealthy, and un-labelled tissue. Cortex specific quantification was performed by masking out subcortical structures from the calculated percentages. Cell counts were performed using the Cell Counter function in FIJI. The VS110 slide scanner depth of field estimation for volumetric cell labelling density was performed by taking a multi-focal-plane image of labelled tissue, finding cells of interest, and moving through the stack to estimate the range by which those cells would be included in cell counting.

#### Electrophysiology

For data recorded from Marmoset A and K, electrophysiological data were recorded using the OpenEphys software and were spike sorted after common mode average referencing and high pass filtering (passband +300 Hz) using Kilosort2^88^. Outputs from the spike sorting algorithms were manually labeled using the ‘phy’ GUI (https://github/kwikteam/phy). Units were accepted if they demonstrated bi-phasic waveforms localized to adjacent channels, had PCA representations showing clear separation from noise, and demonstrated physiologically plausible inter-spike intervals. For data recorded from mice, electrophysiological data were recorded using the Intan RHD2000 recording interface and preprocessed using common mode averaging and bandpass filtering (800-1600 Hz). Custom Matlab scripts identified spikes via user defined amplitude and inter-spike interval criteria for each recording channel. Identified spike waveforms were visualized and channels were rejected if dominated by non-physiological waveforms. Spikes identified using this method were not necessarily well isolated and reflect multi-unit activity present on each recording channel. Local field potentials (LFPs) were averaged for each channel after applying a lowpass filter (passband -300 Hz), and a specialized notch filter for 60 Hz line noise. LFP grand means were calculated from channels that did not show signal inversions or signal drop-offs. This amounted to 48 of the total 64 channels for the plotted means in Figures 3 and 6.

Laser-triggered optogenetic activity was summarized for each sorted unit or recording channel via raster plots and mean firing activity calculated in 1 ms bins from a sliding gaussian window (3 ms for shorter ChR2 stimulation pulses and 15 ms for longer Jaws stimulation pulses). The optogenetic modulation index was calculated at each millisecond bin for each unit by the difference of laser modulated firing rate from a mean baseline firing rate divided by the sum of laser modulated firing rate and mean baseline firing rate. The mean baseline firing rate was calculated using a window of -20 to -5 milliseconds prior to laser onset. Due to the difficulty of measuring inhibitory responses, units in the DLX experiments of Figure 1 were rejected if they had a baseline firing rate of less than 2 Hz. Unit spike waveforms were quantified by the trough to peak duration and also the peak to half height duration. To detect the first instance of laser triggered modulation, it was necessary to first repair an artifactual firing rate depression caused by Kilosort’s removal of spikes coinciding with the laser onset opto-electric artifact. To achieve this, the mean firing rate from 4 ms pre-onset trigger to 4 ms post-onset trigger was spline interpolated using the ‘pchip’ algorithm. Following this, the onset of modulation was calculated as the first instance firing rate became greater or less than baseline plus or minus 2 standard error of the mean (SEM). For the ChR2 and Jaws experiments of Figure 3 and 6, units were rejected from the population of laser modulated units if any point of the baseline firing rate minus 2 SEM and any point of the laser modulation period firing rate minus 2 SEM was less than zero. The ten units plotted in Figure 6J were those units that demonstrated a laser triggered firing rate increase of 1Hz or more from baseline. For these units, a two-sample t-test was used to determine the significance of the difference between ChR2 triggered firing and simultaneous ChR2 + Jaws firing.

## ACKNOWLEDGMENTS

This work is supported by Del Monte Institute for Neuroscience, Schmitt Program on Integrative Neuroscience, and the National Institutes of Health (P50HD103536 to KHW, 5R01EY030998-05 to JFM). We thank members of the Mitchell lab, especially Dina Graf, for technical assistance, and members of the Wang lab for their discussions and technical support. We also thank the Horwitz lab for their thoughtful feedback on the manuscript.

## DECLARATION OF GENERATIVE AI AND AI-ASSISTED TECHNOLOGIES IN THE WRITING PROCESS

The authors declare no use of generative AI or large language models in the production of this manuscript.

## SUPPLEMENTAL FIGURES

**Supplemental Figure 1.**
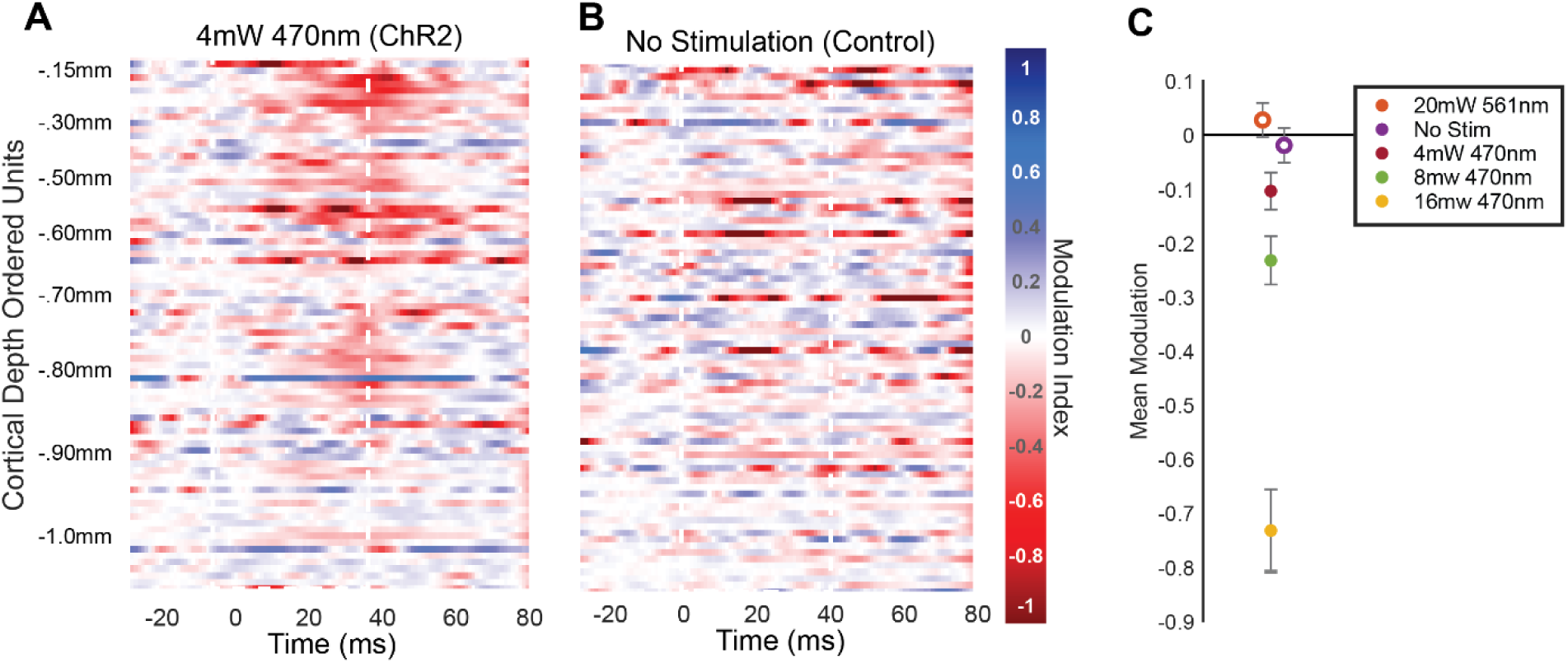
Controlling Laser Power Effects in Awake mDlx-ChR2 Optogenetic Studies in the Marmoset. **A** Modulation scores of units ordered by cortical depth in an additional 4 mW laser power condition, complementing those shown in **Figure 1 L-N**. **B “**No stimulation”-triggered events were randomly selected from periods between runs of laser stimulation. **C** Mean modulation scores were calculated for each condition over the population of units during the laser stimulation window. Error bars indicate +/- 2 SEM. Closed circles indicate p<.05 from a simple one sample students t-test. See **Figure 1 L-N**.

**Supplemental Figure 2.**
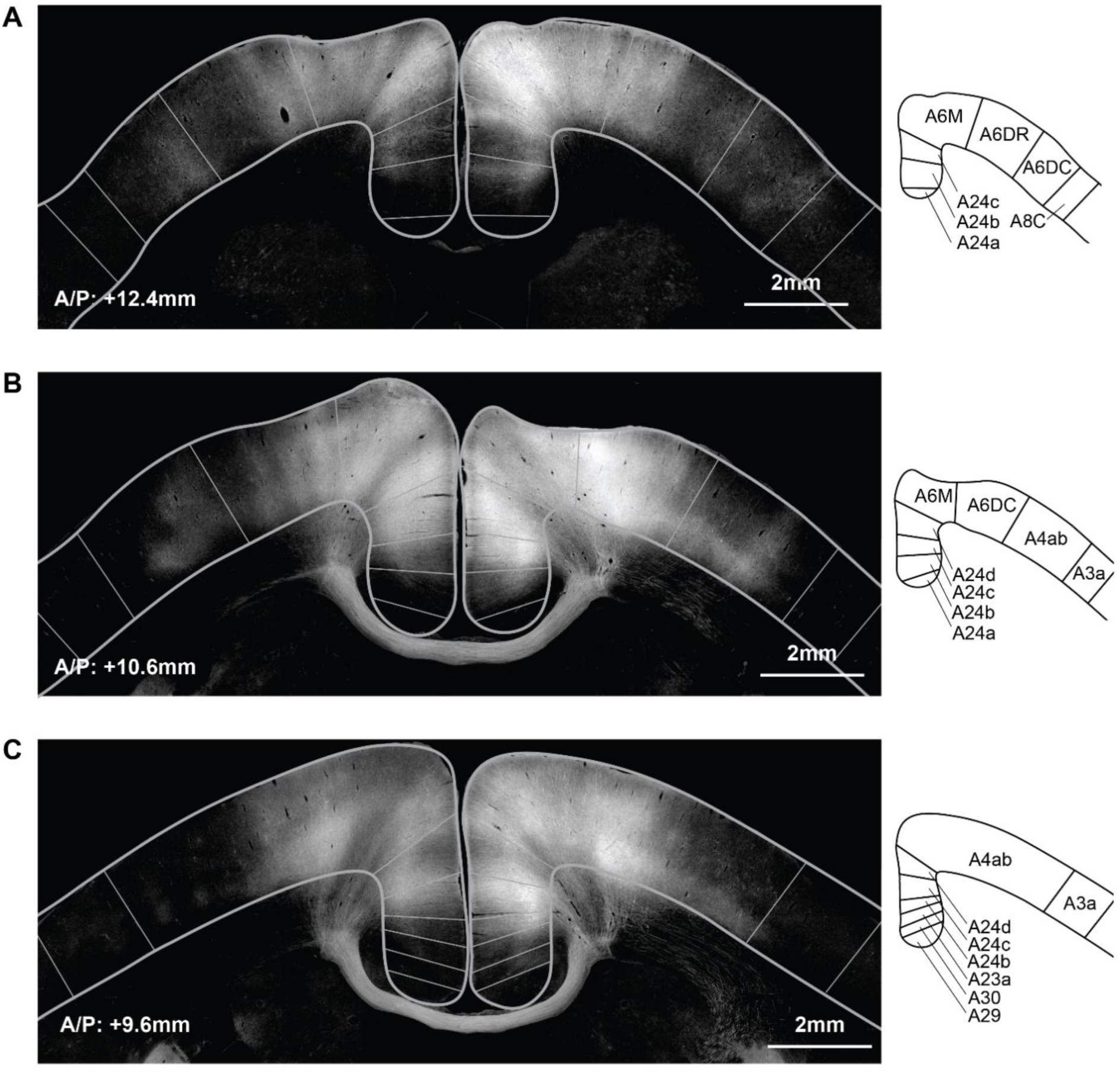
Symmetrical and Patchy Cortical Expression in Cross-Callosal Intersectional Expression of ChR2 Using AAV9. **A-C** Intersectional ChR2 expression in three coronal sections sampled along the anterior/posterior axis. AAVretro-hSyn-Cre was injected into the left cortex, and AAV9-CAG-FLEx-ChR2 was injected into the right cortex, as shown in **Figure 2**. Grey outlines correspond to illustrations in the right column, which are adapted from the Paxino’s Marmoset Brain Atlas (2012). Expression patterns in each section are unevenly distributed, with patchy vertical columns of expression striating the cortex and exhibiting symmetry across the midline. This may reflect multiple adjacent injections or underlying patterns of cross-hemispheric interconnectivity. In **B** and **C**, labeling in the right cortex extends more prominently to deep layers than to superficial layers.

**Supplemental Figure 3.**
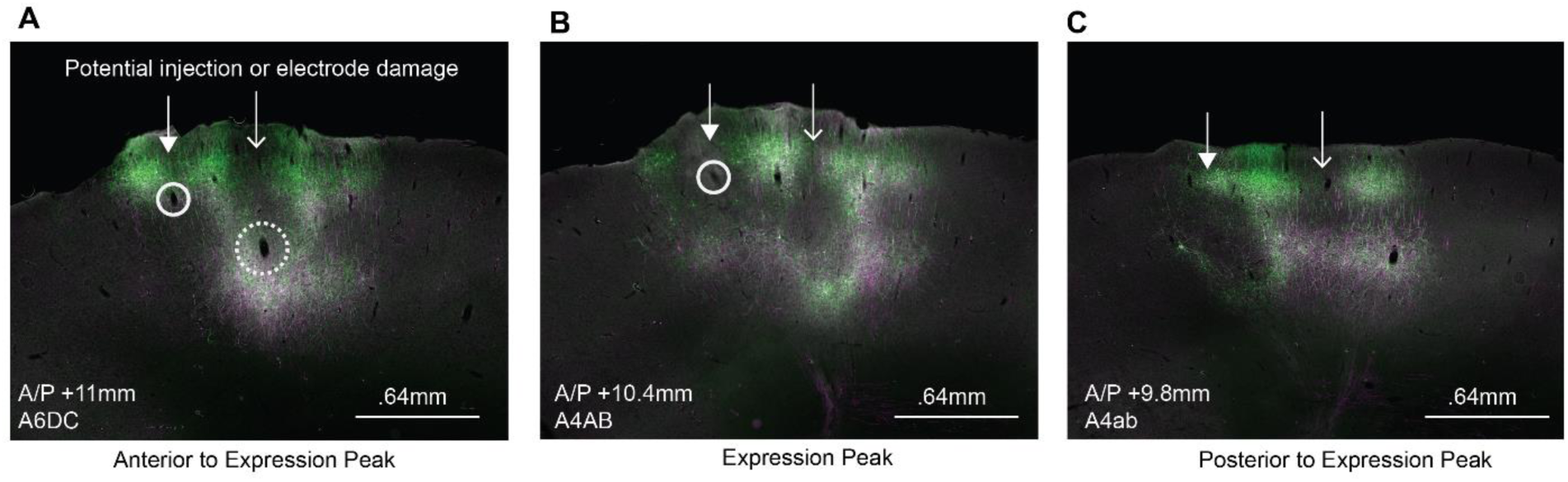
Complex Expression Patterns in Intersectional Cross-Callosal Expression Using AAV8. **A-C** Sequential coronal sections from the cross-callosal AAV8 intersection experiment, showing correspondence between expression shadows (arrows) and underlying vasculature (circles), which could have been ruptured by electrode shanks or the injection pipette. See **Figure 4C**.

**Supplemental Figure 4.**
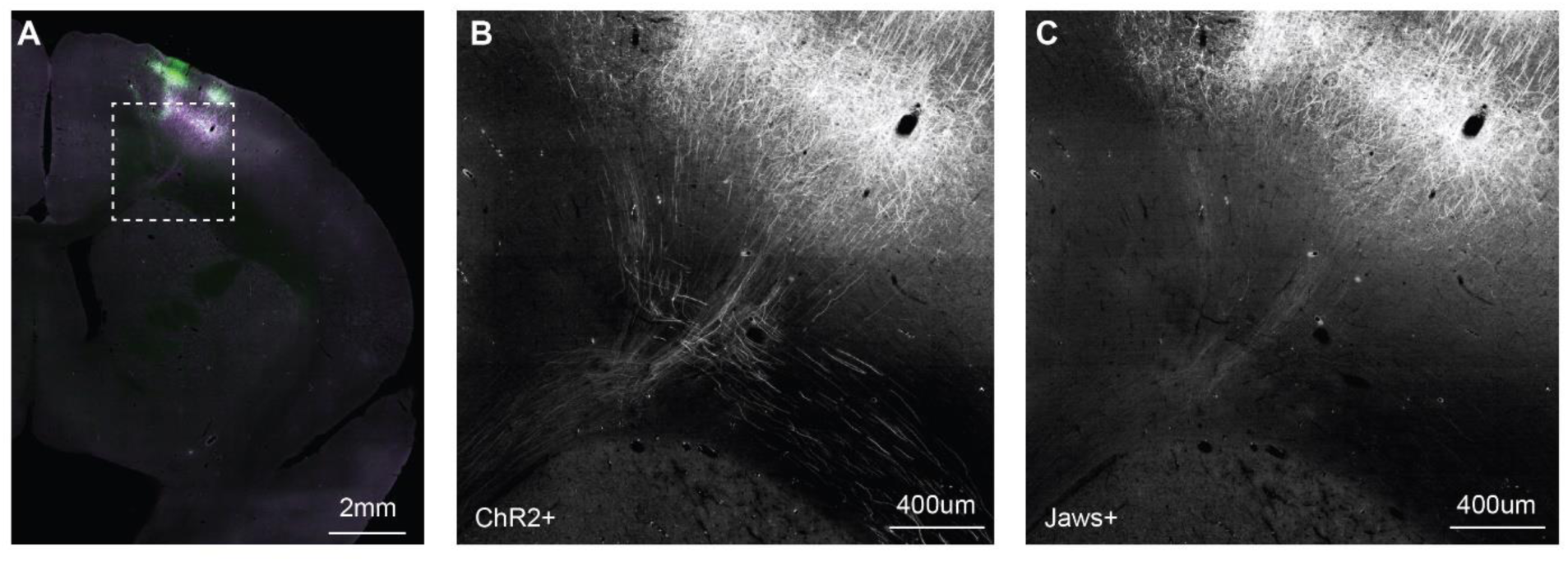
ChR2 Demonstrates More Robust Axonal Labeling than Jaws. **A** Overview of a coronal section labeled with ChR2 and Jaws. The dashed box, which contains deep layers of A4ab, lateral corpus callosum, and dorsal internal capsule, is separated into color channels in **B** and **C**. **B** ChR2 shows robust axonal labeling in projections from the injection site. **C** Jaws shows weaker axonal labeling in comparison to ChR2. See **Figure 4**.

**Supplemental Figure 5.**
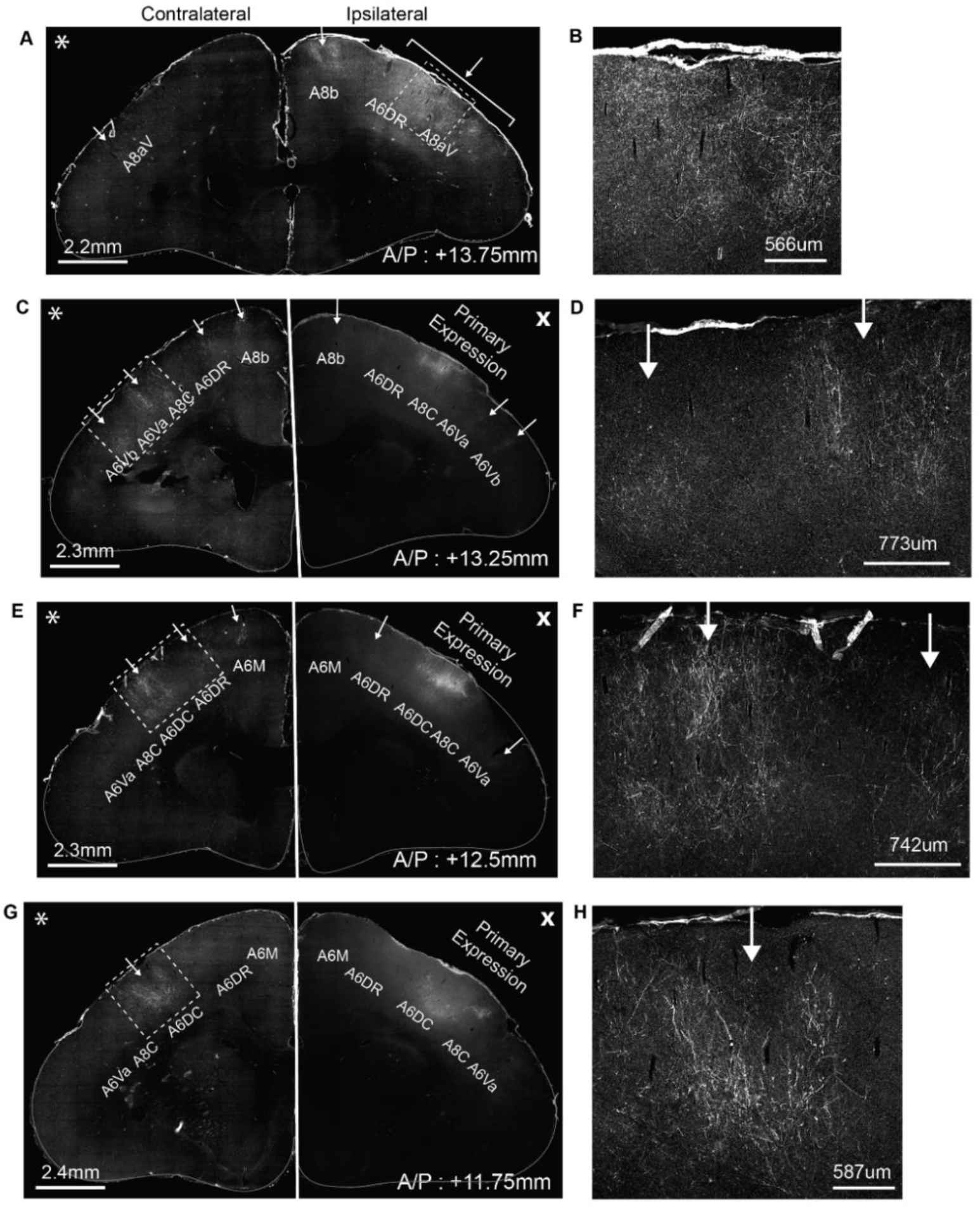
Ipsilateral Premotor-Parietal Projection Neurons Send Frontal and Contralateral Collaterals. All images show ChR2 labeling enhanced with anti-RFP antibody from the premotor-parietal intersection illustrated in **Figure 5**. The right column shows zoom-in views of the dashed rectangles in the left column overviews. Arrows indicate clusters of axon terminals. Panels **C, E, G** are split and adjusted with different imaging parameters to display low-intensity terminal labeling (*) while avoid saturation of high-intensity primary expression (x). **A-B** A section anterior to the injection site shows ipsilateral terminals and a small patch of contralateral terminals in A8aV. **C-D** Contralateral terminal patches are evident in A6a/b, A8C, A6DR, and A8b. **E-F** Contralateral terminal patches are evident in A6DR, A6DC, and A6M. **G-H** Contralateral terminals are evident in a main patch in A6DC.

**Supplemental Figure 6.**
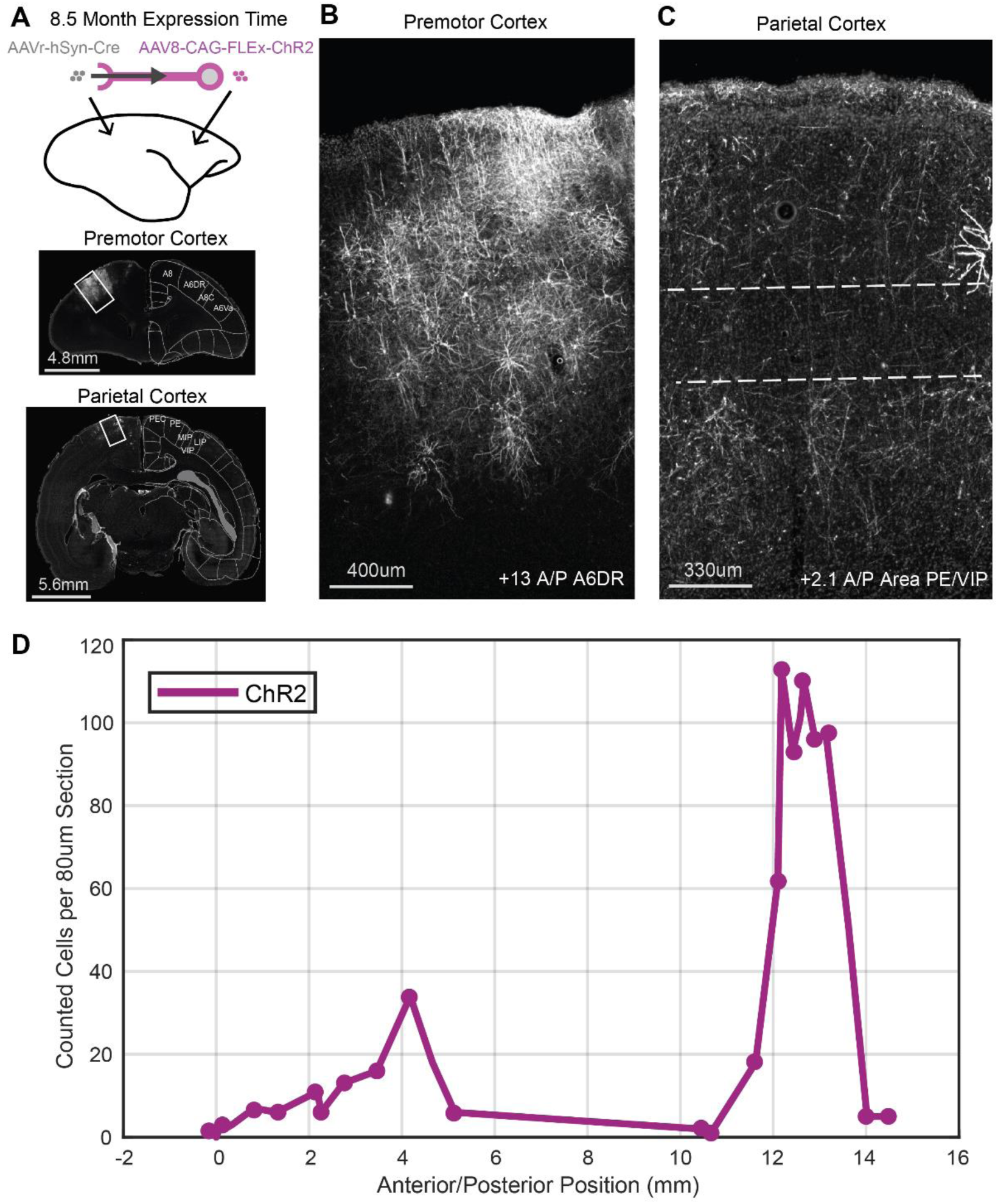
Long Duration Expression of ChR2 in Intersectionally Labeled Premotor-Parietal Project Neurons Using AAV8. **A** Methods for this experiment are the same as described in **Figure 5**, except that ChR2 was injected without Jaws. After 8.5 months of expression, histology and manual cell counting were performed. Full cell labeling is evident without signs of over-expression. Axonal labeling is visible in both deep and superficial layers of the parietal cortex. **B** The injection site in premotor cortex (A6DR) shows robust soma and dendrite labeling. **C** The parietal injection site (PE/VIP) shows dense axonal labeling in superficial and deep layers, with sparse labeling in Layer IV (dashed lines). **D** Manual counting of labelled somas shows greater evidence of leaky expression in the parietal cortex compared to **Figure 5**, possibly due to the longer expression period.

**Supplemental Figure 7.**
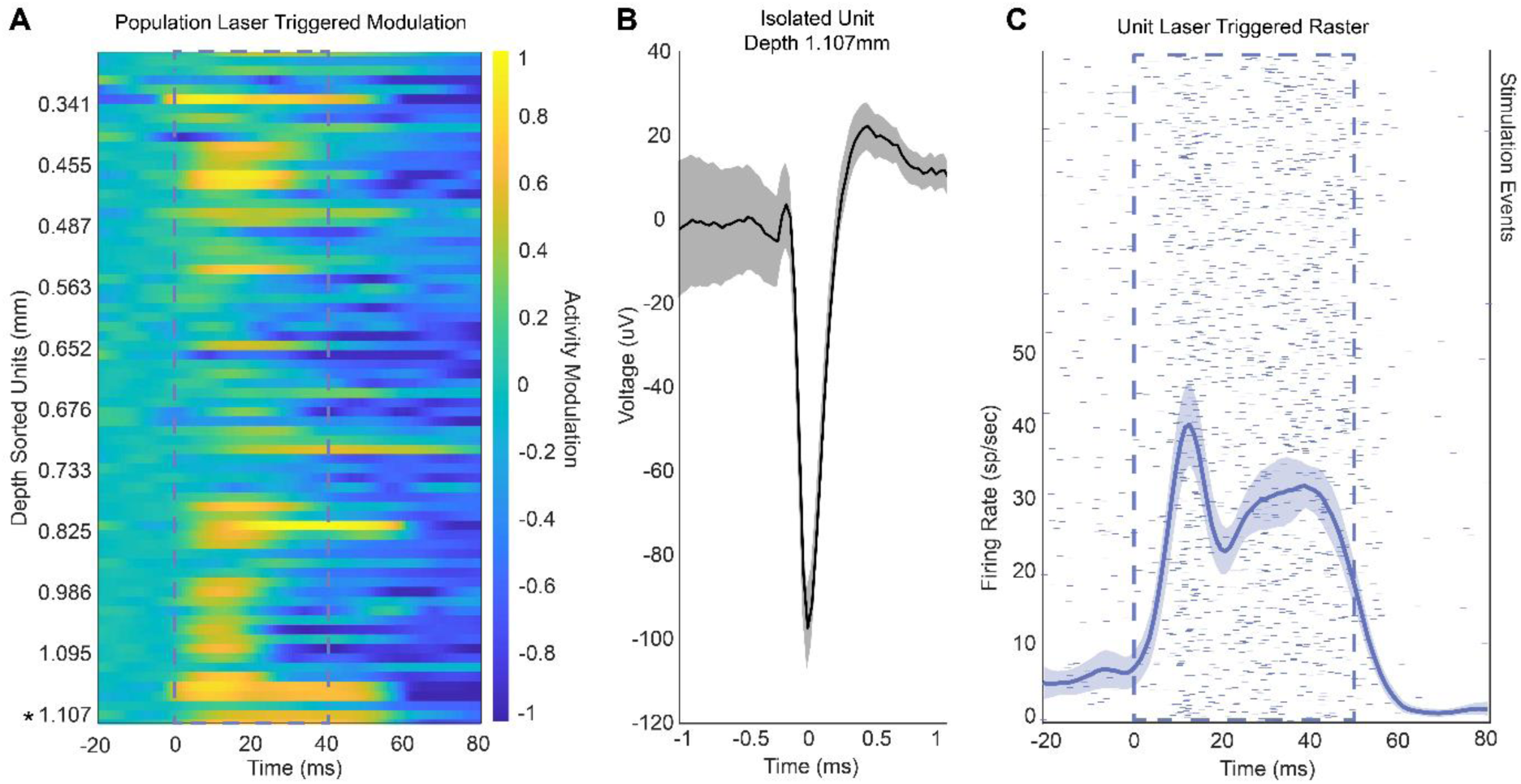
Depth Profile of Intersection-Labeled Units Modulated by ChR2 Stimulation. **A** The modulation of firing rate in response to ChR2 stimulation (yellow=excitatation, blue=suppression) is shown across a population of units, ordered by depth relative to the top of the electrode array, which was inserted fully into the cortex. Data are from a separate electrode insertion and neural population recorded during the experiment in **Figure 6**. The spiking waveform (**B**) and light-triggered firing rate response (**C**) of the deepest unit, found at 1.107 mm (*), are illustrated. Shaded areas indicate 2 SEM. This demonstrates that blue light delivered at the cortical surface can evoke excitation at depths of up to 1.1 mm.

